# Extracellular vesicles from a novel chordoma cell line, ARF-8, promote tumorigenic microenvironmental changes when incubated with the parental cells and with human osteoblasts

**DOI:** 10.1101/2024.10.06.616873

**Authors:** Khoa N Nguyen, Arin N Graner, Anthony R Fringuello, Zoe Zizzo, Lorena Valenzuela, Kamara Anyanwu, Kevin O Lillehei, A Samy Youssef, Samuel Guzman, Christina Coughlan, Michael W Graner

## Abstract

Chordomas are rare, generally slow-growing spinal tumors that nonetheless exhibit progressive characteristics over time, leading to malignant phenotypes and high recurrence rates, despite maximal therapeutic interventions. The tumors are notoriously resistant to therapies and are often in locations that make gross total resections difficult. Here, we describe a new chordoma cell line (ARF-8) derived from an extensive clival chordoma that extended back to the cervical spine. From the cultured cell line we characterized the ARF-8 cellular and extracellular vesicle (EV) proteomes, as well as the impacts of ARF-8 EVs on proteomes and secretomes of recipient cells (both ARF-8 and human osteoblasts) in autocrine and paracrine settings. All the characteristics associated with chordomas as cancers – migration and invasion, therapeutic resistance, metastatic potential – can be driven by tumor EVs. Our proteomic analyses suggested roles for transforming growth factor beta (TGFB) and cell-matrix interactions involving the epithelial-to-mesenchymal transition (EMT), and cell/extracellular matrix interactions in cell migration, consistent with a metastatic tumor phenotype. Our results demonstrated that ARF-8 tumor cell migration was dependent on general (arginine-glycine-aspartic acid [RGD]-based) integrin activity, and ARF-8 EVs could promote such migration. ARF-8 EVs also prompted proteomic/secretomic changes in human osteoblast cells, again with indications that cell-cell and cell-extracellular matrix interactions would be activated. Overall, the EVs promoted predicted tumorigenic phenotypes in recipient cells.

## INTRODUCTION

Chordomas are extremely rare spinal tumors of the sarcoma family with incidences of 0.08 per 100,000^1–4^, or ∼300 cases per year in the US – literally a one-in-a-million disease. The World Health Organization classifies chordomas^5^ as notochordal tumors (referring to their bone sites of origin)^2,6,7^. Spine locations range from clival/skull base (30%-35%), mobile spine (15%-33%), or sacrococcygeal (25%-50%)^1,8^. These slow-growing tumors may feign indolence, but are nonetheless invasively destructive of bone, are notoriously resistant to chemo/radiotherapies, recur frequently without difficult *en bloc* surgical resections, and are often metastatic to other sites (spine and other organs^9,10^). Overall mean survival is 6.29 years; more than 67% of patients survive five years, dropping to less than 40% at 10 years, and approximately 13% at 20 years^1^. Thus, there is significant malignancy associated with chordomas, with no major breakthroughs in treatment for this orphan disease.

Presumably, the best outcomes for chordomas come from *en bloc* or gross total resections (GTRs) accompanied by radiotherapy. These procedures leave clean surgical margins, but anatomical considerations make GTRs difficult^11,12^. Subtotal resections (STRs) are far more prevalent, with inadvertent surgical cellular “seeding” regarded as a potential source of recurrent tumor cells in STRs^13^. Despite the perceived GTRs combined with radiotherapy, 5-and 10-year survival rates only marginally improved compared to STRs and radiation. Tumor cell invasiveness leads to recurrence, so an improved understanding of chordoma migration into and beyond the tumor microenvironment (TME) may help stifle metastatic propensity and improve survival.

All the characteristics typically associated with chordomas as cancers – migration and invasion, therapeutic resistance, metastatic potential – can be driven by tumor extracellular vesicles (EVs)^14,15^. EVs are virus-sized membrane-enclosed nanoparticles released by all cell types^16^, and with profound influences in CNS tumors^17^. EVs are also known to promote bone tumor (osteosarcoma) proliferation, migration, and epithelial-to-mesenchymal transition (EMT)^18^. Their biogenesis includes budding from the plasma membrane or release of multivesicular body contents into the extracellular space^19^. Other extracellular entities include non-membranous nanoparticles such as exomeres and supermeres^20^. EVs are understudied in chordomas with only one published report on them^21^. Herein we describe a novel chordoma cell line, “ARF-8”, characterize its EVs, and study the effects of those EVs on ARF-8 cells and human osteoblast cells (hOBs) as recipients. We determine that ARF-8 EVs dramatically alter the proteomes and secretomes of recipient cells, with implications for the TME and involvement of integrins in ARF-8 cell migration.

## MATERIALS AND METHODS

### Human subjects

Institutional Review Board Statement:

The study was conducted in accordance with the Declaration of Helsinki, and approved by the Colorado Multiple Institutional Review Board (COMIRB) of University of Colorado Anschutz Medical Campus (protocol COMIRB #13–3007) with no expiration date.

Informed Consent Statement:

Informed consent was obtained from all subjects involved in the study. Written informed consent has been obtained from the patient(s) to publish this paper if applicable.

Tissue samples were collected from patients consented to the protocol prior to resection for tumor resection surgeries in the University of Colorado Hospital/Anschutz School of Medicine Department of Neurosurgery. The tissue samples were processed and stored at the Nervous System Biorepository at the Department of Neurosurgery, University of Colorado Anschutz Medical Campus.

### Case Report

The patient was a 22-year-old male with a previous pediatric history of a nonaggressive-appearing mass between the inferior clivus and anterior arch of C1; at the time the leading diagnosis was chordoma vs benign notochordal remnant. A surveillance MRI three months later showed no change in the 1.6 cm infraclival mass. Years later, the patient presented with tongue deviation, nausea, dizziness, and a history of neck pain. MRI (**Fig 1A**) and CT revealed an extensive skull base chordoma extending from the oropharynx to the craniocervical junction. The patient underwent a staged resection, first via an endoscopic endonasal approach, with the second stage as a lateral transcondylar approach with a suboccipital craniotomy and a C1 hemilaminectomy. We obtained tissue from the second surgery, resected in piecemeal fashion, which was 4.0 x 3.0 x 0.5 cm in aggregate, and Pathology diagnosed it as a clinically-recurrent chordoma based on previous determinations and histologic examination of this specimen (**Fig 1B**). The patient received titanium hardware due to the damage at C1-C2; the tumor had encased the vertebral artery and left notable bone destruction, including to the clivus. The patient was also treated for CSF leak and a polymicrobial cervical soft tissue infection. Less than three years later the patient underwent another surgery at an outside hospital after an undisclosed chemotherapy regimen and proton therapy. The patient received further radiotherapy thereafter. The patient is currently alive, with latest imaging showing shrinkage of the residual tumor and no evidence of progression or metastasis.

**Figure 1.**
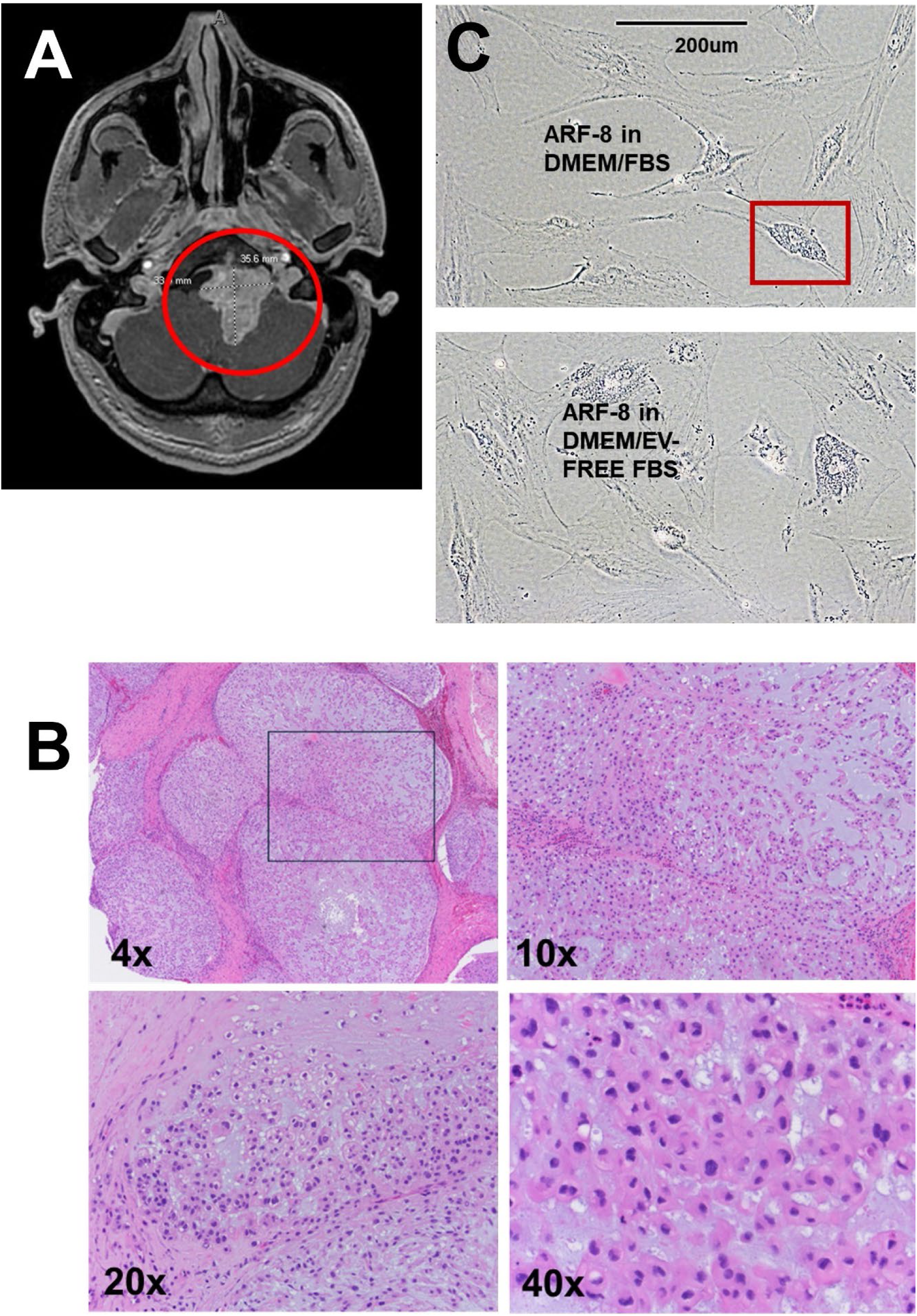
ARF-8 surgical resection, immunohistochemistry, and cell line. A: MRI with a red circle indicating the chordoma in the patient clivus. B: Histological appearance of the chordoma with increasing magnification (4x, 10x, 20x, 40x). The inset from the 4x is shown at higher power in the 10x. The 20x and 40x images are from different slides. C: Appearance of the ARF-8 cell line grown in DMEM+10% fetal bovine serum (FBS; top) and in DMEM+10% EV-free FBS (bottom). Blocked area shows a cell with physaliphorous (bubbly) cytoplasm characteristic of chordomas.

### Cell lines

“ARF-8” cell line was generated from a clival chordoma resected in the Dept of Neurosurgery, University of Colorado Anschutz Medical Campus under COMIRB Protocol 13-3007. The tumor was collected in a sterile container with HGPSA medium (Hibernate A, 1X GlutaMax (Sigma-Aldrich, St Louis, MO, USA), PenStrep, and Amphotericin B (all the rest from Thermo Fisher, Waltham MA, USA). Tissue was diced into 1 mm pieces with scalpels in a tissue culture hood. Tissue pieces were further disaggregated with the blunt end of a 3 ml syringe plunger, and materials were centrifuged at 300 x g for 10 minutes to recover cell material. This was resuspended in 10 ml DMEM medium (Gibco Life Technologies/Thermo Fisher, Waltham MA, USA) + sodium pyruvate (Sigma) supplemented with 10% fetal bovine serum (FBS, Thermo Fisher) and plated in a T-75 flask (Thermo Fisher). Cells were collected and resuspended in DMEM/10%FBS every five days until they finally began to adhere after 4 weeks. Following that, cells were passaged every 4-8 weeks, reaching passage 6 after nearly a year. Cell passage entailed removal of spent media, washing with calcium-and magnesium-free phosphate buffered saline (PBS, Gibco), and treating with 0.25% trypsin (Sigma) for 5 min. Protease activity was stopped by adding 2 volumes DMEM/10%FBS and centrifugal collection of cells (300 x g, 5 min). Cells would then be expanded into new flasks. In the ensuing two years the cells were passaged every 2-4 weeks if in use. Doubling time was determined by cell counting using the following formula: Doubling Time = [T (days) × (ln2)] / [ln (end cell number / beginning cell number)]. Doubling time was ∼19 days (18.8 days, growth rate μ = 0.04).

Human osteoblasts (hOBs) were purchased from PromoCell (C-12720; New York, New York, USA) and cultured in T-75 flasks (Thermo Fisher) in Osteoblast Growth Medium (C-27001) with passage following trypsinization with the Detach Kit (C-41200).

## Extracellular vesicle isolation and characterization

For EV collection, DMEM medium was supplemented with sodium pyruvate and 10% EV-depleted FBS (System Biosciences/SBI, Palo Alto, CA, USA) and cultured for 72 hr intervals before media harvest. We cultured 10 M cells in T-175 flasks (Thermo Fisher) in 15 ml media (12 flasks) until ∼500 ml conditioned media was obtained. EVs were then isolated from conditioned media via differential centrifugation, ultrafiltration/concentration, and ultracentrifugation, as described^22,23^, followed by size-exclusion chromatography (Capto Core 700 resin, Cytiva, Marlborough, MA, USA). Fractions were concentrated using Amicon Ultra-4 10 kDa cutoff centrifugal concentrators (MilliporeSigma, Burlington, MA, USA). EVs were characterized using NanoSight Nanoparticle Tracking Analysis (NTA), Exo-Check arrays, and transmission electron microscopy (TEM).

### NanoSight nanoparticle tracking analysis

We diluted EVs (1:100 or 1:1000) in particle-free phosphate buffered saline (PBS, Gibco) and examined EV numbers and diameters using a NanoSight NS300 (Malvern Panalytical Ltd., Malvern, UK). We used the following analytical settings: camera type: sCMOS, level 11 (NTA 3.0 levels); green laser; slide setting and gain: 600, 300; shutter: 15 ms; histogram lower limit: 0; upper limit: 8235; frame rate: 24.9825 fps; syringe pump speed: 25 arbitrary units; detection threshold: 7; max. jump mode: 5; max. jump distance: 15.0482; blur and minimum track length: auto; first frame: 0; total frames analyzed: 749; temp: 21.099-22.015°C; and viscosity: 1.05 cP.

### Exo-Check arrays

We used Exo-Check antibody-capture arrays (EXORAY200B, System Biosciences) for semi-quantitative detection of 8 putative exosome/EV marker according to the manufacturer’s protocol.

### ExoView

We utilized the ExoView R100 (Nanoview/Unchained Labs, Pleasanton, CA, USA) to characterize and quantify EV sizes and numbers by interferometric imaging, and EV surface markers by immunofluorescence. EVs were analyzed using Tetraspanin kits (Unchained Labs: 251-1044), according to manufacturer’s instructions. In summary, these chips capture EVs based on the three most commonly reported exosome/EV tetraspanin molecules (CD63, CD81 and CD9) and an additional background control, murine IgG. The captured EVs can be subsequently stained for CD63, CD81 and CD9 to determine co-localization of tetrapanins on various EVs. The data were calculated as particles per ml and analyzed for vesicle diameter.

### Transmission electron microscopy

Negative staining of EV samples and observation by transmission electron microscopy (TEM) were performed at the Electron Microscopy Center of the University of Colorado School of Medicine. The EV specimens (5 µl) were dropped onto a 200–400 mesh carbon/formvar coated grid and allowed to adsorb to the formvar (60 sec). Uranyl acetate (2%, 10 µl) was placed on the grid (60 sec). Samples were rinsed in PBS (60 sec). The sample was observed under a transmission electron microscope (FEI Tecnai G2 Biotwin TEM, Thermo Fisher, Waltham, MA, USA) at 80 KV with magnifications of 30,000 to 120,000x. The images were taken by the AMT camera system.

### Brachyury/TBXT detection

#### Western blot

We probed ARF-8 cell lysates (obtained by scraping cells in RIPA buffer (Radio-Immune Precipitation Buffer, R0278; Sigma-Aldrich) supplemented with protease inhibitors and phosphatase inhibitors (cOmplete™ Mini, ethylenediaminetetraacetic acid (EDTA)-free Protease Inhibitor Cocktail, 4693159001; PhosSTOP™, 4906837001; Roche via Sigma-Aldrich). ARF-8 conditioned medium (on cells for 72 hrs) was fractionated as we would obtain from an EV/extracellular particle preparation (12,000 x g pellet; 120,000 x g pellet; 226,000 x g pellet). Equal volumes of sample and 2X Laemmli sample buffer (Bio Rad, cat # 1610737; Bio Rad Laboratories, Hercules, CA, USA) plus 2-mercaptoethanol (50 µl per 950 µl sample buffer, cat # 1610710, Bio Rad) were mixed to generate reducing and denaturing conditions. Samples (15 μg for lysate, as much volume as possible for medium fraction pellets) were electrophoresed on Criterion™ TGX™ (Tris-Glycine eXtended) precast gels (10% Midi gels, cat # 5671033, Bio Rad) with Tris/Glycine/SDS running buffer (Bio Rad, cat # 5671093). Gels were electrotransferred with an iBlot 2 Dry Blotting System (cat # IB21001, ThermoFisher) onto nitrocellulose transfer stacks (cat # IB23001, ThermoFisher). Blots were rinsed in MilliQ water and stained with Ponceau S (cat # P7170, Sigma-Aldrich) to verify transfer and equal loading. Blots were rinsed in Tris-buffered saline (TBS, cat #1706435, Bio Rad) with 0.1% (vol/vol) Tween® 20 (Polysorbate 20) (cat #1706531, Bio Rad; buffer is called TBST). Blots were blocked with 5% non-fat dry milk (NFDM, Nestle Carnation code #1706531; Nestle USA, 28901 NE Carnation Farm Rd Carnation, WA, USA) (wt/vol) in TBST for 1 hr at room temperature on a rocking platform. Blots were rinsed 3 times (5 min per wash) with TBST at room temperature while rocking. Blots were then incubated overnight at 4°C (while rocking) in primary antibody diluted in NFDM 1:1000 (anti-Brachyury/TBXT rabbit monoclonal, clone D2Z3J; Cell Signaling cat # 81694S, Cell Signaling Technologies, Danvers, MA, USA). Blots were washed in TBST as described above. Secondary antibody was a goat anti-rabbit (H+L; Cell Signaling, cat #7074P2) IgG with HRP label, used at 1:1000 dilution in 5% NFDM/TBST. It was incubated on the blots on a rocker for 1 hr at room temperature. Following washings as described, blots were treated with SuperSignal™ West Femto Substrate developer (cat #34096; ThermoFisher) and bands were visualized using an iBright FL1500 device (Invitrogen/Thermo Fisher, A44241, Carlsbad, CA, USA).

### Extracellular vesicle incubation with recipient cells

For EV treatment experiments for proteomic studies, ARF-8 cells were cultured in T-75 flasks (1M cells/flask) in DMEM supplemented with sodium pyruvate and 10% EV-depleted FBS as above. hOB cells were grown in T-75 flasks (1 M cells/flask) in Osteoblast Growth Medium as above. Prior to treatment with EVs, media were removed, and cells were washed with PBS. EVs were diluted directly to 100 μg/ml in culture medium and filtered through a sterile 0.45 μm syringe filter before application to the recipient cells for 24 hrs. After 24 hrs, media were harvested and centrifuged (2000 x g, 10 min, RT) and then frozen for later experiments. Cells were rinsed three times with PBS, and were scraped directly off the plates in residual PBS. Cells were pelleted in a microfuge (17K x g, 20 min, 4°C), PBS was aspirated, and cell pellets were flash-frozen in an ethanol/dry ice bath and stored at -80°C until use. For proteomic studies, EVs were isolated as described above, and were flash-frozen before storage at -80°C until use.

### Mass spectrometry-based proteomic analysis

Samples were handled by the Mass Spectrometry Proteomics Shared Resource Facility on the CU Anschutz Campus.

### Sample digestion

The samples were digested according to the FASP (Filter-Aided Sample Preparation) protocol using a 10 kDa molecular weight cutoff filter. In brief, the samples were mixed in the filter unit with 200 µL of 8 M urea, 0.1M ammonium bicarbonate (AB) pH 8.0, and centrifuged at 14,000 g for 15 min. The proteins were reduced with 10 mM DTT for 30 min at room temperature (RT), centrifuged, and alkylated with 55 mM iodoacetamide for 30 min at RT in the dark. Following centrifugation, samples were washed 3 times with urea solution, and 3 times with 50 mM AB. Protein digestion was carried out with sequencing grade modified Trypsin (Promega) at 1/50 protease/protein (wt/wt) at 37°C overnight. Peptides were recovered from the filter using 50 mM AB. Samples were dried in Speed-Vac and desalted and concentrated on Thermo Scientific Pierce C18 Tip.

### Mass spectrometry analysis

Samples were analyzed on an Orbitrap Fusion mass spectrometer (ThermoFisher) coupled to an Easy-nLC 1200 system (ThermoFisher) through a nanoelectrospray ion source. Peptides were separated on a self-made C18 analytical column (100 µm internal diameter, x 20 cm length) packed with 2.7 µm Cortecs particles. After equilibration with 3 µL 5% acetonitrile 0.1% formic acid, the peptides were separated by a 120 min linear gradient from 6% to 38% acetonitrile with 0.1% formic acid at 400 nl/min. LC mobile phase solvents and sample dilutions used 0.1% formic acid in water (Buffer A) and 0.1% formic acid in 80% acetonitrile (Buffer B) (Optima™ LC/MS, ThermoFisher). Data acquisition was performed using the instrument supplied Xcalibur™ (version 4.1) software. Survey scans covering the mass range of 350–1800 were performed in the Orbitrap by scanning from m/z 300-1800 with a resolution of 120,000 (at m/z 200), an S-Lens RF Level of 30%, a maximum injection time of 50 milliseconds, and an automatic gain control (AGC) target value of 4e5. For MS2 scan triggering, monoisotopic precursor selection was enabled, charge state filtering was limited to 2–7, an intensity threshold of 2e4 was employed, and dynamic exclusion of previously selected masses was enabled for 45 seconds with a tolerance of 10 ppm. MS2 scans were acquired in the Orbitrap mode with a maximum injection time of 35 milliseconds, quadrupole isolation, an isolation window of 1.6 m/z, HCD collision energy of 30%, and an AGC target value of 5e4.

### Protein identification and databases

MS/MS spectra were extracted from raw data files and converted into mgf files using Proteome Discoverer Software (ver. 2.1.0.62). These mgf files were then independently searched against the human database using an in-house Mascot server (Version 2.6, Matrix Science). Mass tolerances were +/- 10 ppm for MS peaks, and +/- 25 ppm for MS/MS fragment ions. Trypsin specificity was used allowing for 1 missed cleavage. Met oxidation, protein N-terminal acetylation, and peptide N-terminal pyroglutamic acid formation was allowed as variable modifications. Scaffold (version 5.0, Proteome Software) was used to validate MS/MS-based peptide and protein identifications. Peptide identifications were accepted if they could be established at greater than 95.0% probability as specified by the Peptide Prophet algorithm. Protein identifications were accepted if they could be established at greater than 99.0% probability and contained at least two identified unique peptides. Volcano plots were generated in GraphPad Prism 9.0 using Scaffold data. Orthogonal partial least squares-discriminant analysis (PLS-DA), Variable Importance in Projection (VIP) scores, and hierarchical clustering heatmaps from ANOVA statistical analyses were generated using the MetaboAnalyst 6.0 online platform (https://www.metaboanalyst.ca/MetaboAnalyst/ModuleView.xhtml) with sum and range scaling normalizations. Proteomic data were further analyzed using QIAGEN Ingenuity Pathway Analysis (IPA; QIAGEN Inc., Hilden, Germany https://digitalinsights.qiagen.com/IPA). We also used FunRich Functional Enrichment Analysis (http://www.funrich.org/) for some analyses.

### Cytokine release/‘secretome’ assays

Following treatment with ARF-8 EVs (24hrs), the conditioned media from either ARF-8 cells or from hOBs were assayed with Proteome Profiler Human XL Cytokine Arrays (ARY022B; R&D Systems, Minneapolis, MN, USA) performed as per manufacturer’s protocols. Arrays were developed with chemiluminescence (SuperSignal™ West Femto Substrate, 34096; ThermoFisher) and spot intensities quantified using an Alpha Innotech FluorChem Q System (ProteinSimple/Bio-Techne). Background was subtracted from each membrane, and average luminosity from duplicate spots was compared between treatment groups. Results were displayed as heatmaps using Microsoft Excel (Microsoft 365; Microsoft, Redmond, WA, USA).

### Protease arrays

Conditioned media was collected from EV-treated or untreated (control) astrocytes and handled as described above. We incubated 1.5 ml of media on Protease Arrays (R&D Systems Proteome Profiler Human Protease Array Kit, Cat # ARY021B; R&D Systems Inc., 614 McKinley Place NE, Minneapolis, MN 55413, USA), and followed the manufacturer’s instructions. Arrays were developed, quantified, and displayed as described above.

### Protease assays

Matrix metalloproteinase (MMP) activities (as a general enzyme class) were assayed using abcam MMP Activity Assay Kits (Cat # ab112146, Fluorescent [green]; abcam, 152 Grove Street, Suite 1100 Waltham, MA 02453, USA) following manufacturer’s instructions. Samples were ARF- 8 EVs directly (50μg/ml). Measurements were performed under kinetic conditions for 0-1 hr on a plate reader (excitation/emission = 490/525 nm; Biotek Synergy H4 hybrid reader, Winooski, VT, USA, software version Gen5 3.09).

### Migration/scratch assays

Migration capacity of ARF-8 cells exposed to 0, 50, or 100 µg/ml ARF-8 EVs was measured using standard procedures for a scratch assay, essentially as described^22,24^. Briefly, ARF-8 cells at near confluency in 6-well plates (Corning) were scratched with a 200 µl pipet tip to generate a gap between cells. ARF-8 EVs were added (or not), and scratch-gap closure was measured at 16 or 24 h (compared to the distance at 0 h). Measurements of ‘gap closure’ were performed using NIH Image J. Fetal bovine serum (10%) was added to medium as a positive control attractant; medium with EV-free FBS was the baseline attractant. For some experiments, RGD peptide at 1 μM (GRGDNP; Cat. No.: HY-P1740; MedChemExpress, Monmouth Junction, NJ, USA) was used to inhibit ARF-8 cell migration measured at 16 hrs.

### TGFB/TGFβ (transforming growth factor β) detection and neutralization/inhibition

#### Western blot

We probed fresh DMEM/10%FBS medium (never on cells), ARF-8 conditioned medium (on cells for 72 hrs) and fractions of the conditioned medium as we obtain from an EV/extracellular particle preparation (12,000 x g pellet/supernatant; media filtered through 0.22 μm filters and concentrated from 500 ml down to 24 ml; 120,000 x g pellet/supernatant from that; 226,000 x g pellet/supernatant from the 120K x g supernatant; 270,000 x g pellet/supernatant from the 226K x g supernatant). Western blot procedure was followed as above. Fractions (30 μg) were electrophoresed on Criterion™ TGX™ (Tris-Glycine eXtended) precast gels (4-20% Midi gels, cat # 5671033, Bio Rad) with Tris/Glycine/SDS running buffer (Bio Rad, cat # 5671093). Gels were transferred to nitrocellulose blots as described above. Blots were then incubated overnight at 4°C (while rocking) in primary antibody at 0.1 μg/ml (chicken IgY anti-TGFB1/1.2; biotechne/R&D systems Catalog #: AF-101-NA). Blots were washed in TBST as described above. Secondary antibody was a goat anti-chicken IgY (H+L; Thermo Fisher, A16054) IgG with HRP label, used at 1:5000 dilution in 5% NFDM/TBST. It was incubated on the blots on a rocker for 1 hr at room temperature. Following washings as described, blots were treated with SuperSignal™ West Femto Substrate developer (cat #34096; ThermoFisher) and bands were visualized using an iBright FL1500 device (Invitrogen/Thermo Fisher, A44241, Carlsbad, CA, USA).

### Neutralization/inhibition

To test effects of blocking TGFB on ARF-8 cells or their EVs, we treated ARF-8 cells or EVs with TGF beta-1,2,3 antibody (MA5-23795; clone 1D11; Thermo Fisher) at 1 μg/ml. This was done directly onto cells in 6-well plates (500,000 cells/well) in medium for 2 hrs prior to replacement with fresh medium for 24 hrs. EVs were treated likewise, and then separated from free antibody by centrifugation at 120K x g for 1 hr. Pellets were resuspended in fresh DMEM/10%FBS and incubated with ARF-8 cells for 24 hrs. Isotype control antibody was murine IgG1 control 13C4 (Genzyme (Sanofi), Cambridge, MA, USA) used at the same concentration. After media removal, cells were washed 3 times with PBS, and subjected to a CellTiter-Glo assay.

### CellTiter-Glo 2.0 assay

CellTiter-Glo 2.0 Luminescence Assay (Promega, Madison, WI, USA) was used for ATP detection according to the manufacturer’s directions. Briefly, after PBS rinses, cells were treated with 1 ml CellTiter-Glo 2.0 reagent and placed on an orbital shaker for 2 min, then left at RT for 10 min. Cells were then pipetted out and distributed into an opaque, clear-bottom 96-well plate (100 μl/well; greiner Bio One, Monroe, NC, USA; item 655090-99) and luminescence detected with a Bio Tek Synergy H4 hybrid reader (software version Gen5 3.09; Agilent Technologies, Englewood, CO, USA).

### Data analysis

Data were statistically assessed using Microsoft Excel and/or GraphPad Prism software (GraphPad Prism 9.5, San Diego, CA, USA). Student’s t-test or one-way or two-way ANOVA analyses were performed depending on experimental circumstances. ANOVAs were followed by pairwise multiple comparisons with Tukey’s method to show the significant differences between treatments and groups. Data were presented as mean ± SEM (or SD); significance was set at 0.05. In some cases, internal algorithm statistics were applied by the web-based programs.

## RESULTS

ARF-8 was a clinically recurrent, extensive clival chordoma that extended into the cervical spine (C1). The patient had a pediatric history of a much smaller chordoma or benign notochord remnant. The recurrence covered a much greater area and exhibited bone destruction while enclosing the vertebral artery. The tumor was removed in two stages via a transsphenoidal approach followed by a lateral craniotomy. **Fig 1A** shows an MRI with a “grapelike cluster” in the clivus, and histologic appearances typical of chordomas (**Fig 1B**) with the lobular presentation at low magnification of chords and nests of cells in a myxoid matrix with the lobules separated in some cases by thick fibrous bands. Higher magnification views reveal mono-and occasional bi-nucleated cells with vacuolated, physaliphorous cytosols.

ARF-8 cells were cultured in DMEM supplemented with 10% FBS and eventually could be grown in DMEM supplemented with EV-free FBS where they maintained a physaliphorous cytoplasmic appearance (**Fig 1C**). Growth rate μ = 0.04; doubling time was approximately 19 days.

ARF-8 EVs were harvested by standard means from 72 hr conditioned medium by filtration, differential centrifugation, and size exclusion chromatography (**Fig 2A**). We characterized vesicle size and concentration via nanoparticle tracking analysis (NTA, Nanosight) with mean and mode diameters of 195nm and 165nm, respectively, and a typical concentration of ∼1.8 x 10^7^ EVs/ml (**Fig 2B**). Notably, these are nearly ten-fold lower concentrations than we frequently see from other tumor cell types^22^. EV sizes and bilayer membrane phenotypes were validated by transmission electron microscopy (TEM, **Fig 2C**). Additional characterizations came from ExoCheck arrays showing the presence of CD9, CD81, CD63, FLOT1, ICAM1, ALIX, TSG101 as EV markers (**Fig 2D**). Additional verification of membrane markers (CD81, CD63, and CD9, including their colocalization on EVs) as well as sizes and concentrations were obtained by single particle interferometric reflectance imaging sensing (SP-IRIS) and immunofluorescence (ExoView, **Fig 2E**). Of note, SP-IRIS consistently shows smaller particle sizes compared to NTA^25,26^, as the latter’s sizing is based on hydrodynamic radius. The EV counts, based on chip volume and dilution, translate into ∼2-3 x 10^7^ EVs/ml. These characterizations and features indicate that these are “small” EVs^27^. Finally, we note that brachyury/TBXT is detected by Western blot in ARF-8 cell lysates, but also in pellets from the sequentially fractionated conditioned medium (12,000 x g pellet; 120,000 x g pellet; 226,000 x g pellet, **Fig 2F**). While we isolated other extracellular particles for brachyury/TBXT analysis, we note that the bulk of our proteomic analysis and functional impacts will focus on ARF-8 “small EVs” (now called “EVs” throughout).

**Figure 2.**
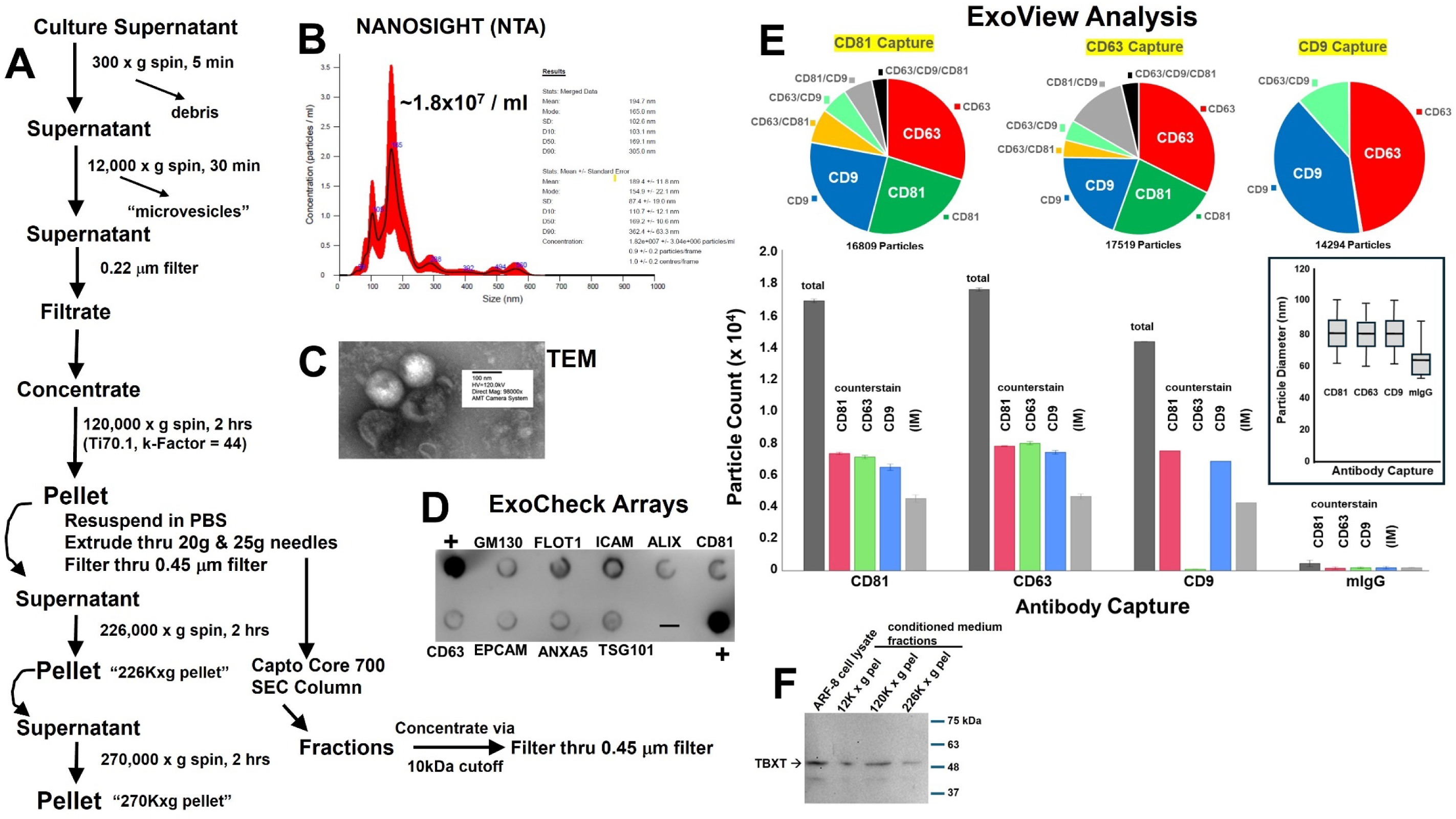
ARF-8 EVs and extracellular particles. A: Schematic of steps involved in EV/particle isolation. B: Nanosight nanoparticle tracking analysis (NTA) of ARF-8 EVs (Stats: Merged Data; Mean: 197.7 nm, Mode: 165.0 nm, SD: 102.6 nm), C: Transmission electron microscopy (TEM) of ARF-8 EVs with scale bars listed. D: ExoCheck arrays showing putative typical EV markers for lysed EVs from the ARF-8 cell line. “+” = positive control. E: ExoView single particle interferometric reflectance imaging sensing (SP-IRIS) with immunofluorescence counterstain; top row shows pie charts of proportions of counterstained CD81, CD63, and CD9 markers on immunocaptured EVs; bottom row shows particle counts by capture and counterstain (including negative control murine IgG); inset to right shows particle diameters based on antibody capture type. F: Western blots of ARF-8 cell lysates and fractionated conditioned media extracellular pellets (12,000 x g; 120,000 x g; 226,000 x g) probed with anti-brachyury/TBXT antibody.

We performed proteomic analyses on the ARF-8 EVs, identifying nearly 300 proteins (**Supp Table 1**). The ExoCarta database lists the “top 100” proteins found in the culmination of their archived entries http://exocarta.org/exosome_markers_new and ARF-8 EVs contain 33 of those (**Fig 3A**), again validating these as bona fide EVs. Using Ingenuity Pathway Analysis (IPA) for gene ontology and pathway derivations, we see over half of ARF-8 EV proteins’ (subcellular) origins are from the plasma membrane and extracellular space (including 14 collagen subunits) with the rest consisting of cytoplasmic and nuclear origin proteins (**Fig 3B**). The highest scoring IPA network is “Connective Tissue Disorders, Dermatological Disease and Conditions, Developmental Disorder”; Score = 39; 23 Focus Molecules, with TGFB1 as a major node (**Fig 3C**). Overlaying the top 7 networks into a radial pattern places transforming growth factor beta (TGFB) as the central node component (**Fig 3D**). IPA Upstream Analysis scores TGFB1 most significantly, and the TGFB Group Mechanistic Network is shown in **Fig 3E**. The top 25 Canonical Pathways extracted from the data show ties to numerous extracellular and intracellular signaling pathways, as well as cell-cell and cell-matrix interaction pathways (**Fig 3F**). IPA Network Legends (node and path design shapes, edges and their descriptions) are in **Supp Fig 1**. The IPA summary for ARF-8 EV proteomics is in **Supp Fig 2.** FunRich^28^ fundamental enrichment analysis top Biological Pathways pointed to numerous integrin-related pathways as well as epithelial-to-mesenchymal transition (EMT; **Fig 3G**). We point out that ARF-8 EVs harbor five integrin subunits (ITGB1, ITGA11, ITGAV, ITGA2, and ITGBL1). We believe this is the first proteomic analysis performed on chordoma EVs.

**Figure 3.**
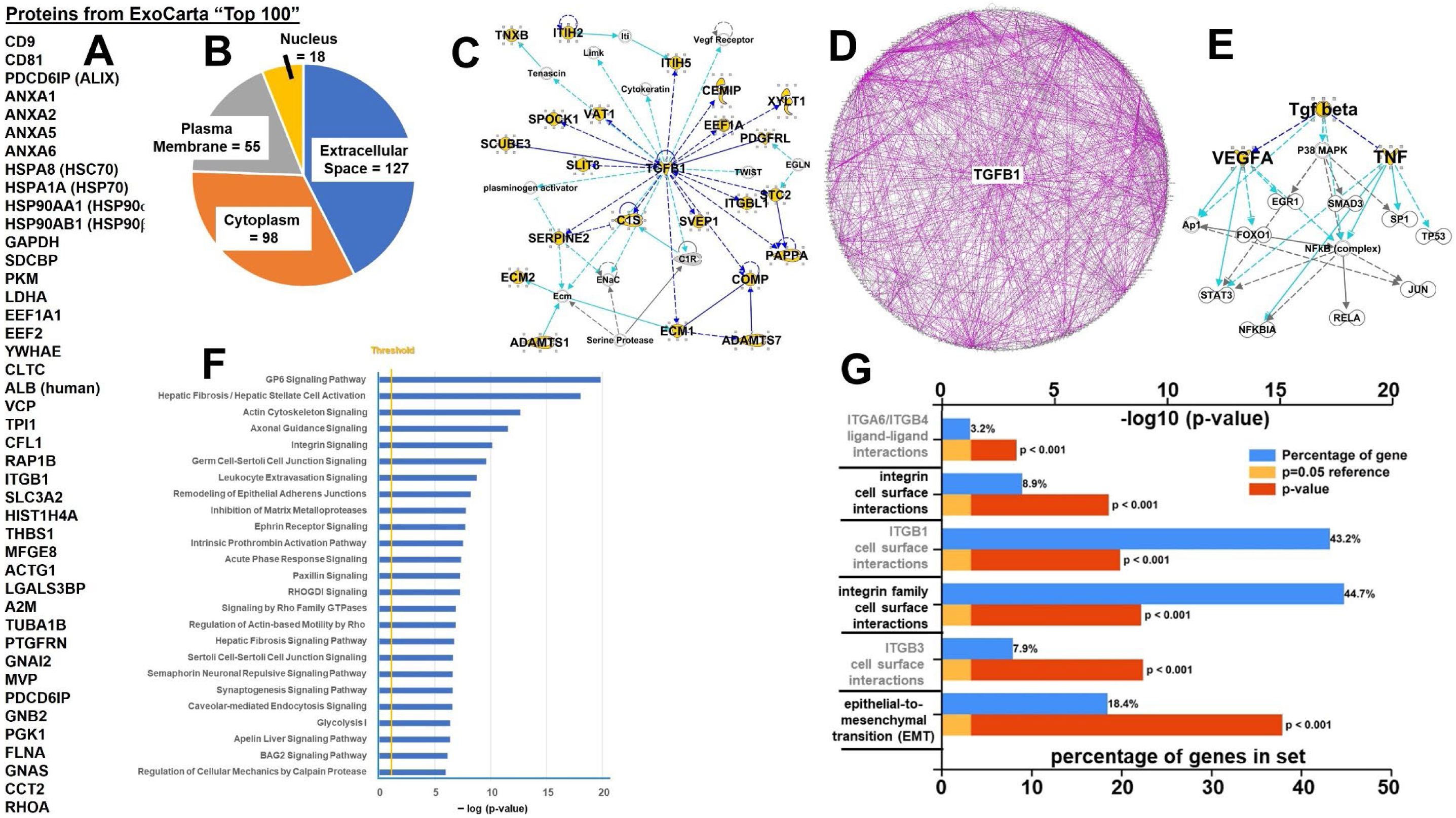
ARF-8 EV proteomics. A: Thirty-three ARF-8 EV proteins that are among the ExoCarta “Top 100” proteins most frequently found in that database (http://exocarta.org/exosome_markers_new). B. Subcellular/extracellular derivation of ARF-8 EV proteins. C. Ingenuity Pathway Analysis (IPA) highest scoring network “Connective Tissue Disorders, Dermatological Disease and Conditions, Developmental Disorder”; Score = 39; 23 Focus Molecules. Note TGFB1 as a major node. Gold highlights = proteins identified in proteomics and secretome analyses. “Scores” are based on Fisher’s exact test, -log(p-value); “Focus Molecules” are considered focal point generators within the network. Number of genes illustrated is limited to 35 by the algorithm. IPA Network Legends (node and path design shapes, edges and their descriptions) are in **Supp Fig 2.** D. Overlay of top 7 IPA-generated networks in radial view; again, TGFB1 is the central node. E. IPA Upstream Analysis scores TGFB1 most significantly, TGFB Group Mechanistic Network is shown. F. IPA-derived Top 25 Canonical Pathways. Statistical significance “Threshold” (gold line) = 1.25. G. FunRich Top Biological Pathways shown by statistical significance and percentage of identified genes in the overall dataset.

We incubated ARF-8 cells with their own EVs for 24 hrs and measured changes in the recipient cell proteomes and secretomes compared to the parental (untreated) cells. We identified over 3300 proteins in untreated ARF-8 cells, and over 3500 proteins in the EV-treated cells (**Supp Table 2**) with an overlap of 3090 proteins (Venn diagram, **Fig 4A**). The Principal Component Analysis Scores Plot indicates clear separation of the replicate datasets (**Fig 4B**), with Variable Importance in Projection (VIP) scores indicating the variables (genes/proteins) driving the PLS-DA results (**Fig 4C**). Volcano plots show the significant changes in protein expression in treated versus control settings (**Fig 4D**), and this is reiterated in the hierarchical clustering heatmaps of the top 100 features (**Fig 4E**). IPA Graphical Summary (summary statement is in **Supp Fig 3**) for the comparison of untreated vs EV-treated cells (**Fig 4F**) emphasizes cell movement, migration, and invasion, along with many other tumorigenic categories (tumor microenvironment, tumor proliferation, angiogenesis). The FunRich top 15 Biological Pathways (**Fig 4G**) compared proteomes of ARF-8 cells untreated and treated with ARF-8 EVs shown as the percentage of genes in the datasets, and showed increased pathway components in many cell-surface/extracellular signaling pathways. Notable among these are proteoglycan signaling, as ARF-8 cells and EVs express syntenin-1/syndecan binding protein/SDCBP; ARF-8 cells express glypicans GPC1, GPC4, and GPC6, while the EVs express GPC1. Integrin signaling is also implicated, with 5 integrin subunits from ARF-8 EVs, and 13 subunits from ARF-8 cells. These results suggest an active extracellular environment surrounds ARF-8 cells, particularly when those cells are incubated with their own EVs.

**Figure 4.**
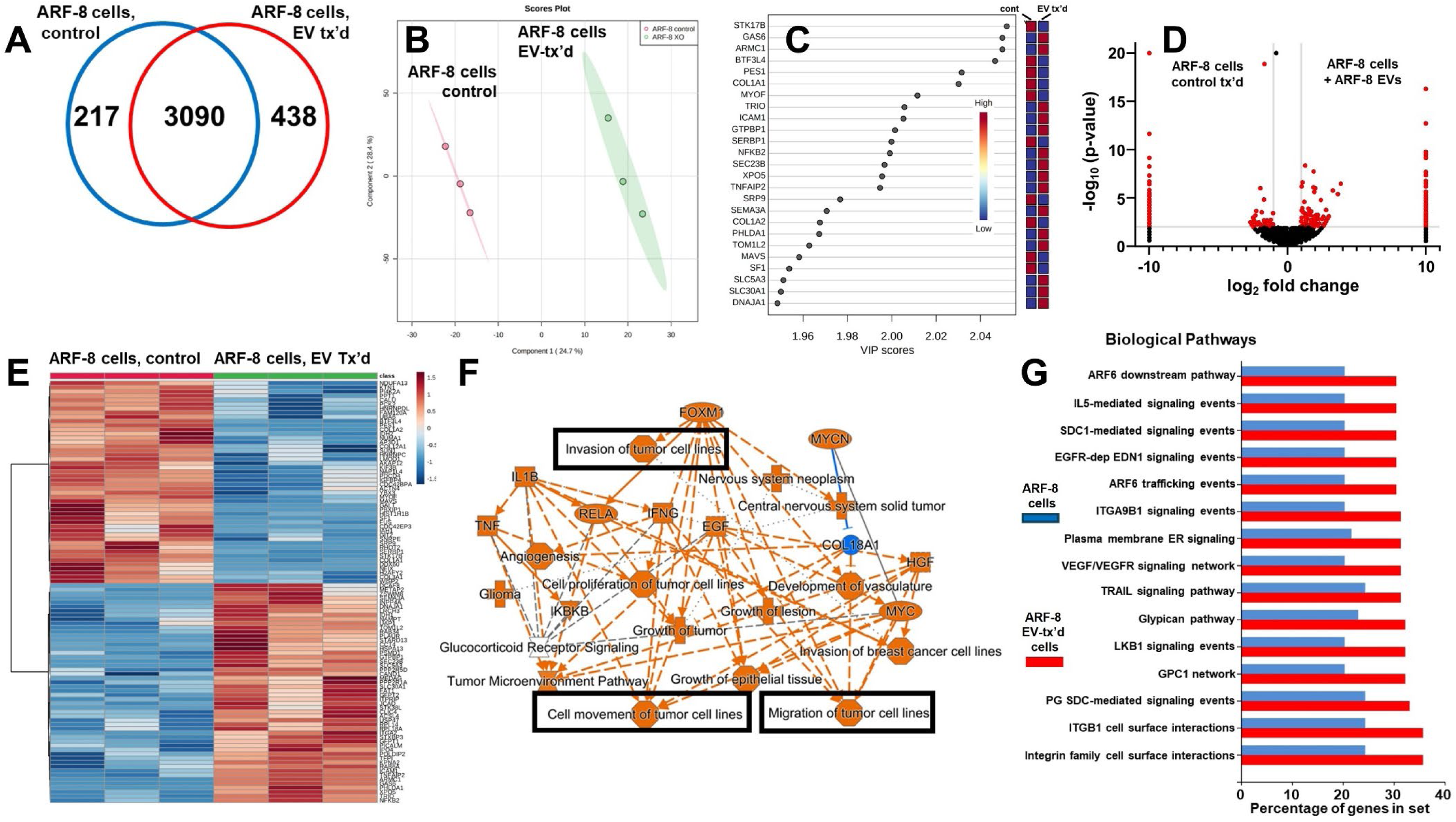
ARF-8 cellular proteomics without and with treatment by ARF-8 EVs. A. Venn diagram showing unique and overlapping numbers of proteins identified. B. Orthogonal partial least squares discriminant analysis (PLS-DA) Score Plot showing clustering and distinction between ARF-8 cell proteomics (controls vs cells treated with ARF-8 EVs). C. Variable Importance in Projection (VIP) scores indicating the variables’ (genes/proteins) driving the PLS-DA results. D. Volcano graphs showing differential proteomic expression between control ARF-8 cells and ARF-8 cells treated with ARF-8 EVs. Data were generated in Scaffold 5 and presented using Prism 9.5. E. Hierarchical clustering heatmaps from ANOVA statistical analyses utilized normalized data that were standardized by autoscaling features (top 100) with Euclidean distance measurements and clustered by ward; Control ARF-8 cells, left; EV-treated ARF-8 cells, right. F. IPA Graphical Summary of the ARF-8 proteomes from untreated vs EV-treated cells; orange colors indicate putatively activated states and pathways, while blue predicts downregulation. Items of particular interest regarding cell motility are boxed in black. G. Top 15 Biological Pathways generated in FunRich identified comparing proteomes of ARF-8 cells (blue bars) vs ARF-8 cells treated with ARF-8 EVs (red bars) shown as the percentage of genes in the datasets. Data for B, C and E were generated in Metaboanalyst 5.0.

ARF-8 cells incubated with their own EVs alter their cyto-chemokine “secretome”, measured here in conditioned media using a 109-component antibody array, **Fig 5A**, as well as their release of proteases (also assessed by an antibody array, **Fig 5B**). We see increases and decreases of both pro-and anti-inflammatory molecules upon treatment with EVs (e.g., increased IL6, IL8, TNFA, but also IL10 and soluble Fas ligand, **Fig 5A**). There are also increases in some secreted proteases (**Fig 5B**), although matrix metalloproteinases (MMPs) 2 and 7 remain high under both conditions. The relationships among some of the secreted factors are illustrated in the IPA-generated interactome (Network #1 “Cellular Movement, Hematological System Development and Function, Immune Cell Trafficking”; Score = 47; 22 Focus Molecules, **Fig 5C**). Notably, increases in CCL7 and sustained high levels of CCL2 could be involved in monocyte attraction^29^ but also in recruitment or conversion of monocytes to myeloid-derived suppressor cells in the TME^30^. Further IPA analysis of Diseases and BioFunctions of the EV-treated cell secretome identified “cell migration” as one of the top categories (**Fig 5D**, among other interesting findings), implying active proteases and cell movement. We directly measured collagenase (MMP) activity of ARF-8 EVs (**Fig 5E**) and further showed that ARF-8 EVs could promote migration of ARF-8 cells in scratch fill assays (**Fig 5F**, with measurements graphed in **Fig 5G**). At 100 μg/ml, ARF-8 EVs significantly promoted migration compared to EV-free medium (purple bar vs gray bar). Thus, predicted cell surface and extracellular activity from the EV and cell proteomics (**Figs 3**, **4**) were validated in functional assays.

**Figure 5.**
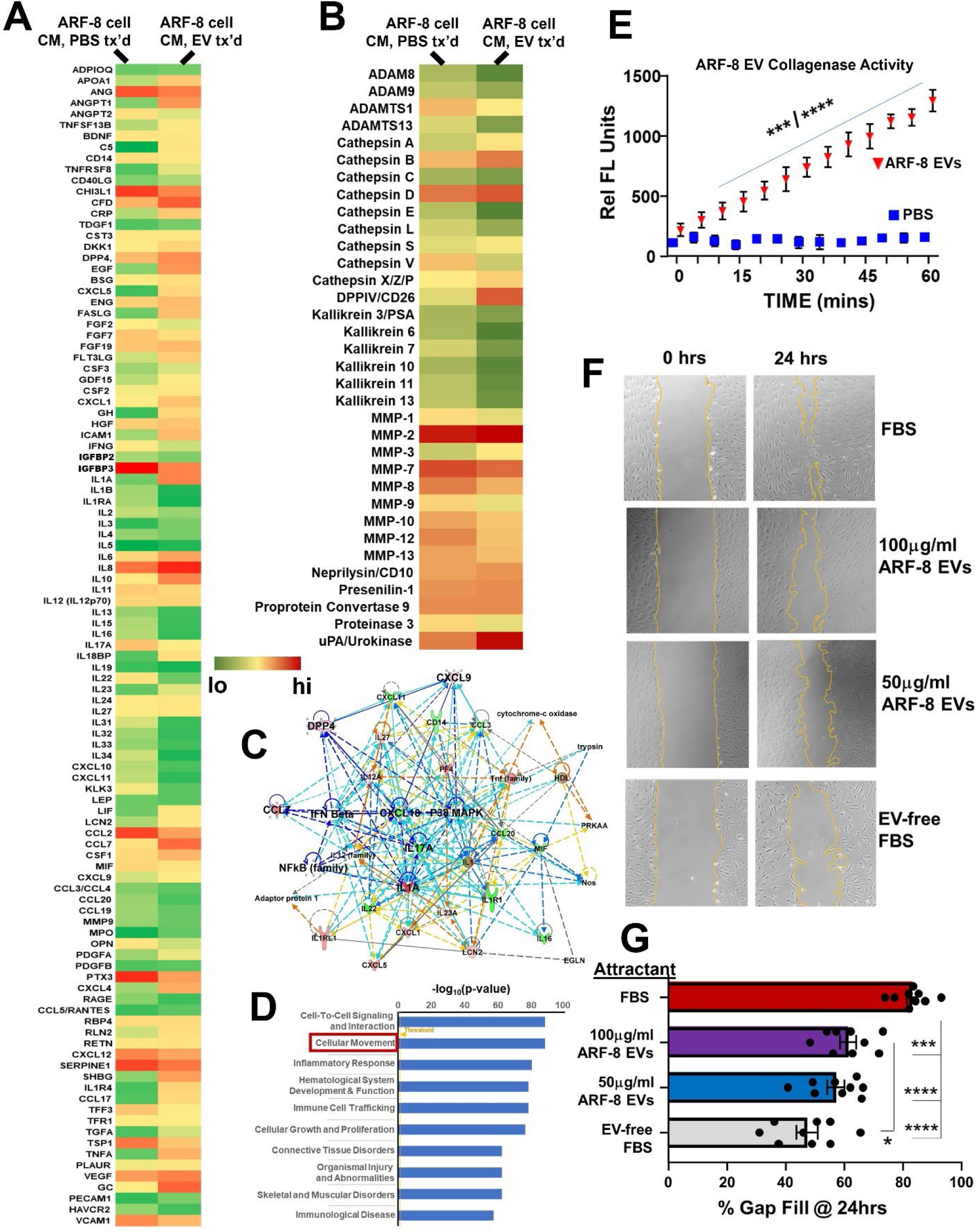
ARF-8 Cell Secretome +/- incubation with ARF-8 EVs. A. Secretome (cyto/chemokine array) heatmap of ARF-8 cell conditioned media (CM) from cells treated (“tx’d”) with PBS or treated with ARF-8 EVs. B. Heatmap of secreted proteases from ARF-8 cells treated with PBS or treated with ARF-8 EVs. C. Ingenuity Pathway Analysis (IPA) Network #1, “Cellular Movement, Hematological System Development and Function, Immune Cell Trafficking”; Score = 47; 22 Focus Molecules. D. ARF-8 secretome top 10 (by statistical significance) Diseases and BioFunctions from IPA. Boxed is “Cellular Movement” as the second highest-scoring category. “Threshold” value for significance is shown in orange = 1.25. E. ARF-8 EV collagenase/MMP activity compared to PBS. *** p<0.001 vs PBS up to 30 min time point; **** p<0.0001 vs PBS for 30-60 min time points. F. Migration/scratch assay of ARF-8 cells filling the gap over 24hrs. Attractants are fetal bovine serum (FBS, + control), ARF-8 EVs at two different concentrations, and EV-free FBS (baseline attractant). G. Quantification of scratch assay. * p<0.05; *** p<0.001; ****p<0.0001.

Further, the proteomic data strongly suggested involvement of integrins (**Figs 3**, **4**) that could be related to cell migration. To test this, we subjected ARF-8 cells to scratch fill assays in the absence or presence of an RGD (arginine-glycine-aspartic acid) peptide inhibitor (GRGDNP). ARF-8 cell proteomics indicate that the cells express at least four possible integrin combinations (ITGAVB1, -B3, -B5, ITGA5B1) that likely bind RGD as a ligand^31^. GRGDNP strongly inhibited cell migration at 1 μM concentration (**Fig 6 A,B**) without resulting in cell death (not shown; treatment did induce notable cell layer detachment in some cases). Log dose increases (10 μM, 100 μM) did result in increasing cell death (not shown).

**Figure 6.**
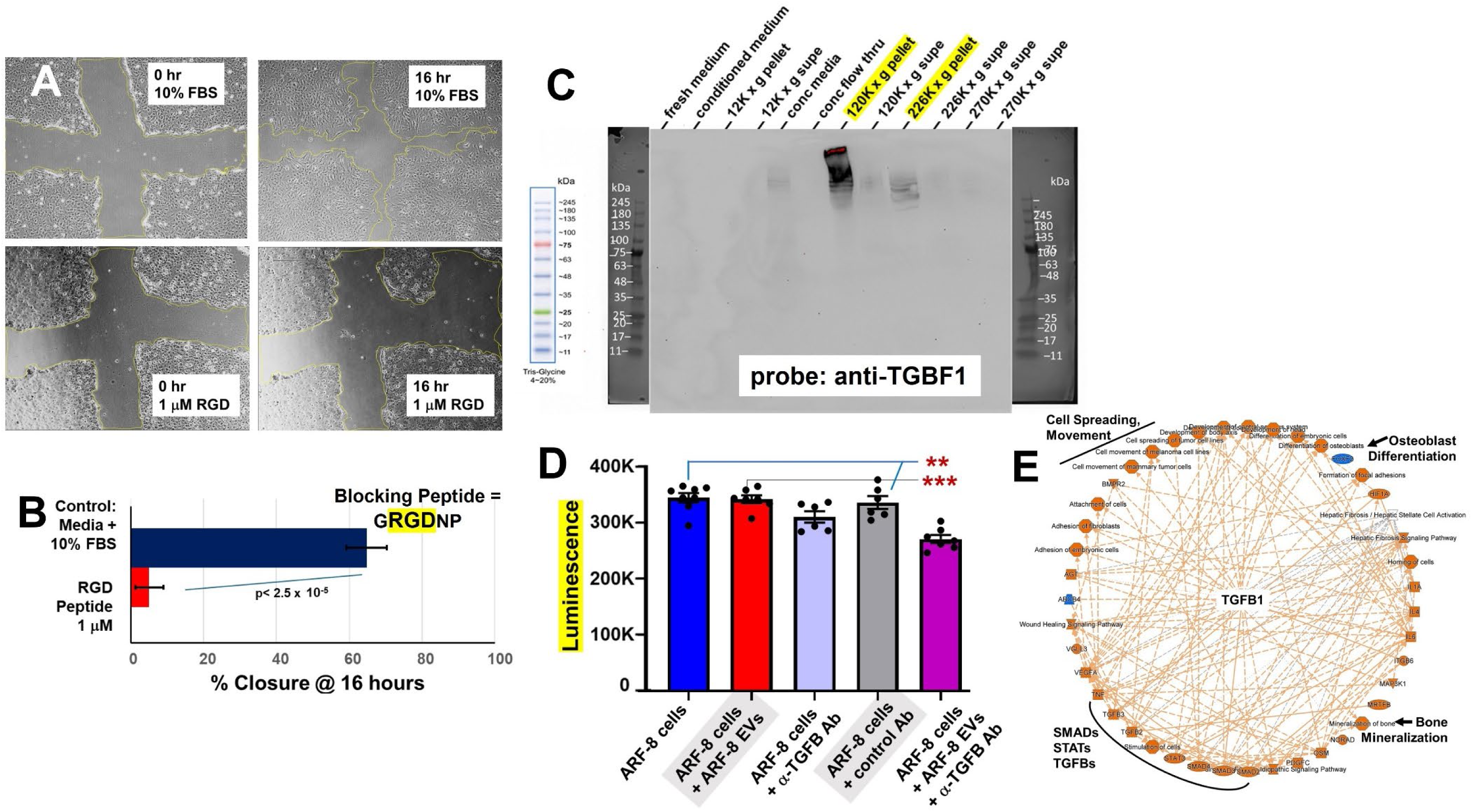
Integrins, TGFB, and EMT in ARF-8 cells. A. Scratch closure assays with ARF-8 cells moving into the gap after 16hrs treatment with either medium + 10% FBS (positive control) or in the presence of 1uM GRGDNP peptide to block integrins. B. Quantification of scratch closure after 16 hrs where RGD peptide strongly inhibited cell migration. C. Western blot probing for TGFB1 of fractionated ARF-8 conditioned medium (30 μg for each fraction) showing reactivity in high-speed centrifuged pellets (marked in yellow, 120,000 x g, and that supernatant further centrifuged at 226,000 x g). Molecular weight markers on the blot are indicated, and shown separately at left. D. Treatment of ARF-8 EVs with anti-TGFB1 antibody 1D11 before EV incubation with ARF-8 cells reduces ATP production. E. ARF-8 EMT signature proteome represented as an IPA Graphical Summary in radial format shows TGFB1 as the major central node with links to SMAD transcription factors, STAT signaling, cell migration, and osteoblast differentiation and bone mineralization upregulated activities.

Transforming growth factor beta (TGFB) was a prominent feature in the pathway analyses of ARF-8 EVs (**Fig 3**), and we identified apparently latent TGFB1 in Western blots of ARF-8 EVs (120K x g pellets, characteristic of small EVs or exosomes) as well as in 226K x g pellets (characteristic of so-called “exomeres”). This indicates that at least the latent form of TGFB1 (>245kDa) is more prevalent in vesicular or particulate forms vs soluble forms (**Fig 6C**). Using a known function-blocking antibody (mAb 1D11), we incubated ARF-8 cells and their EVs with mAb 1D11 and found the blocking TGFB1 on ARF-8 EVs prior to incubation with ARF-8 cells reduced ATP generation by ARF-8 cells (**Fig 6D**). While this effect was statistically significant, it was nonetheless rather small, suggesting that TGFB effects on cell metabolism might not be dramatic.

IPA analysis of ARF-8 EVs (**Fig 3**), found associated molecules and pathways involving TGFB, and FunRich implied that epithelial-to-mesenchymal transition (EMT) pathways may exist, as might be expected for a migratory/metastatic cancer expressing beta-catenin and a Wnt family member^32,33^ along with TGFB and numerous integrins. We mined the “EMTome”, described in^34^, which lists over 1000 genes/proteins/microRNAs that are thought to constitute EMT signatures. Our ARF-8 cellular, secretome, and EV data were compared and trevealed >300 proteins from the ARF-8 proteomes within the EMTome (**Supp Table 3**). Entering those expression values into IPA (summary in **Supp Fig 4**) allowed us to find the Graphical Summary (**Fig 6E**) that, in radial layout, again placed TGFB1 as the central node, with connections to SMAD transcription factors, cell motility, and curiously, osteoblast differentiation and bone mineralization.

The latter findings prompted us to consider osteoblasts as cells of the chordoma TME, in this case, in the clivus, where there was bone degradation due to tumor invasion. Spinal osteoblasts are disorganized in a RAS-driven zebrafish chordoma model^35^, suggesting that such cells may be affected by the chordoma extracellular environment. We tested this by treating human osteoblasts (hOBs) with ARF-8 EVs and conducting proteomic and secretomic assays on the hOBs and their conditioned media. We identified over 3700 proteins in untreated and EV-treated cells (**Supp Table 4**) with an overlap of 3405 proteins (Venn diagram, **Fig 7A**). The Principal Component Analysis Scores Plot indicates clear separation of the replicate datasets (**Fig 7B**), with Variable Importance in Projection (VIP) scores indicating the variables (genes/proteins) driving the PLS-DA results (**Fig 7C**). Volcano plots showed the significant changes in protein expression in treated versus control settings (**Fig 7D**), and this was reiterated in the hierarchical clustering heatmaps of the top 100 features (**Fig 7E**). IPA Graphical Summary (summary statement in **Supp Fig 5**) for the comparison of untreated vs EV-treated cells (**Fig 7F**) emphasized cell migration and invasion, roles for TGFB and integrins, and neoplastic and malignant characteristics. The FunRich top 12 Biological Pathways (**Fig 7G**) compared proteomes of untreated hOBs and those treated with ARF-8 EVs shown as the percentage of genes in the datasets. We see numerous proteoglycan and integrin signaling pathways implied, again suggesting that ARF-8 EVs promoted cell-cell and cell-matrix interactions. hOBs express glypicans GPC1, -4, and -6, syndecan SCD2, and syntenin-1/syndecan binding protein/SDCBP. hOBs also express 13 integrin subunits.

**Figure 7.**
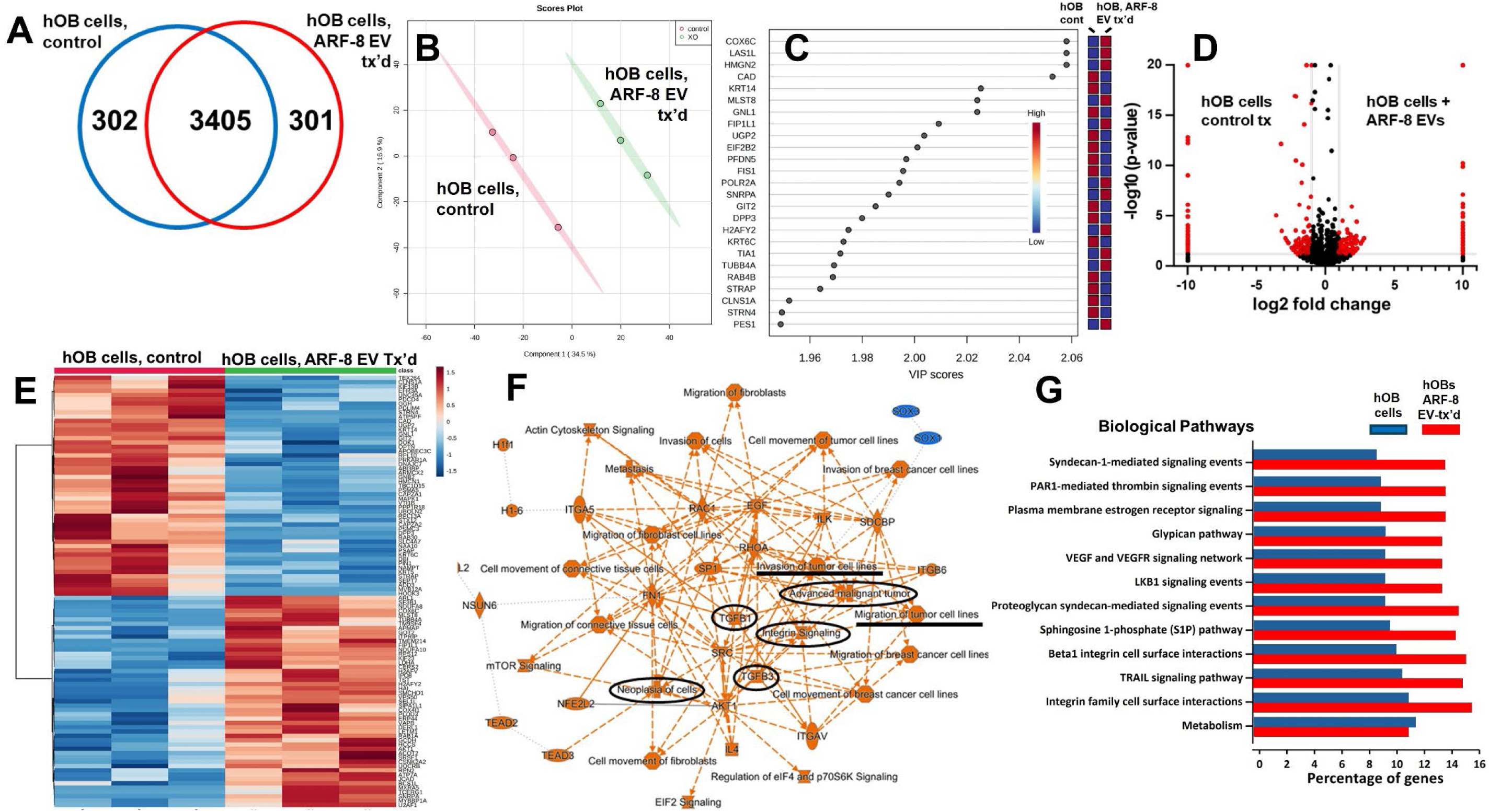
Human osteoblast (hOB) cellular proteomics without (control, PBS-treated) and with ARF-8 EV treatment (tx’d). A. Venn diagram showing unique and overlapping numbers of proteins identified. B. Orthogonal partial least squares discriminant analysis (PLS-DA) Score Plot showing clustering and distinction between hOB cell proteomics (controls vs cells treated with ARF-8 EVs). C. Variable Importance in Projection (VIP) scores indicating the variables’ (genes/proteins) driving the PLS-DA results. D. Volcano graphs showing differential proteomic expression between control hOB cells and hOB cells treated with ARF-8 EVs. Data were generated in Scaffold 5 and presented using Prism 9.5. E. Hierarchical clustering heatmaps from ANOVA statistical analyses utilized normalized data that were standardized by autoscaling features (top 100) with Euclidean distance measurements and clustered by ward; Control hOB cells, left; ARF-8 EV-treated hOB cells, right. F. IPA Graphical Summary of the hOB proteomes from untreated vs EV-treated cells; orange colors indicate putatively activated states and pathways, while blue predicts downregulation. Items of particular interest regarding cell motility, integrins, TGFBs and neoplasia/malignancy are circled or underlined in black. G. Top 12 Biological Pathways generated in FunRich identified comparing proteomes of hOB cells (blue bars) vs hOB cells treated with ARF-8 EVs (red bars) shown as the percentage of genes in the datasets. Data for B, C and E were generated in Metaboanalyst 5.0.

hOBs incubated with ARF-8 EVs also altered their cyto-chemokine “secretome” (measured here in conditioned media as in **Fig 5**). Some changes are notable (**Fig 8A**), such as the increase in BDNF, which is associated with enhanced chondrosarcoma cell motility^36,37^ and angiogenesis via VEGF^38^. Molecular profiling of chordoma cell lines suggested that OPN, upregulated here, may

**Figure 8.**
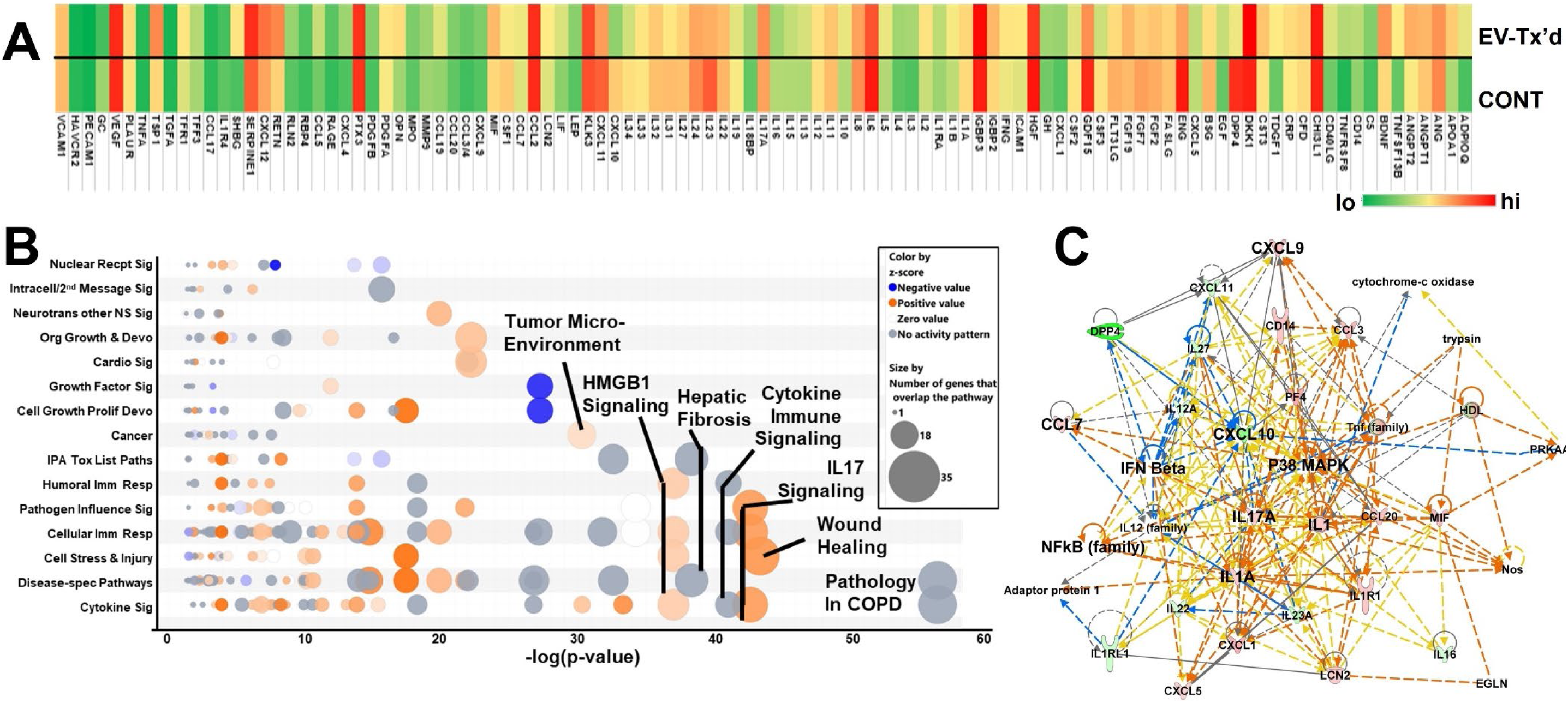
hOB secretome following incubation with ARF-8 EVs. A. Secretome (cyto/chemokine array) heatmap of hOB cell conditioned media (CM) from cells treated (“tx’d”) with PBS or treated with ARF-8 EVs. B. Ingenuity Pathway Analysis (IPA) Canonical Pathways, “Bubble Chart”, generated from the protein quantification changes in the ARF-8 EV-treated hOB secretome (**Fig 8A**). Key pathways are denoted. Significance threshold (-log[p-value]) is set at 1.25. Blue indicates a negative activation z score; orange is positive, gray shows no activity pattern. C. IPA Network #1, “Cellular Movement, Hematological System Development and Function, immune Cell Trafficking”; Score = 47; 22 Focus Molecules.

be part of the tumorigenesis program in chordoma^39^, and is also well known for promoting bone metastases^40^. The thrombospondins, including TSP1, are recognized for the impacts in bone biology, but also for their roles in bone metastasis^41^. We note increased levels of SHBG from ARF-8 EV-treated hOBs, as well. While the data on SHBG in bone cancers and metastasis are sparse, one publication notes that in patients with prostate cancer, higher serum SHBG levels are associated with reduced bone mineral density and osteoporosis^42^.

Using IPA’s Canonical Pathways “Bubble Chart” to display the statistical values of categories and sub-categories of high-scoring pathways of the secretomes (**Fig 8B**), we noted that fibrotic and wound healing pathways, along with activated immune stimulation and “danger signals” (HMBG1) scored highly and appeared activated according to z score; interestingly, “Tumor Microenvironment” activation also appears prominently. The highest-scoring IPA network “Cellular Movement, Hematological System Development and Function, Immune Cell Trafficking” (Score = 47; 22 Focus Molecules) shows potential increased activity of CCL7 with decreases in CXCL10 but sustained high levels of CCL2 (**Fig 8C**) that could be involved in monocyte attraction^29^ but also recruitment or conversion to myeloid-derived suppressor cells in the TME^30^, as noted in **Fig 5A,C**. Increased levels of CCL7, CXCL9 and CXCL10 were found in plasma of breast cancer patients^43^. Osteoblasts downregulate bone angiogenesis via CXCL9 following STAT1 activation^44^, tying bone biology to immune readouts. Overall, the continuing theme is one of extracellular activation upon stimulation by ARF-8 EVs.

## DISCUSSION

Chordomas are rare, generally slow-growing spinal tumors that nonetheless exhibit progressive characteristics over time, leading to malignant phenotypes and high recurrence rates, despite maximal therapeutic interventions^1,2^. As clival chordomas are a subset of these tumors, large studies with consistent treatment attempts followed over time are few (e.g.,^45,46^). Moreover, likely due to their overall rarity, molecular characterizations across chordomas are minimal compared to other cancer types, leading to a paucity of targets and a lack of biological information that could lead to better therapies^47^.

Here, we describe a new chordoma cell line (ARF-8) derived from a chordoma that expanded from an apparent non-aggressive mass or benign notochordal remnant to an extensive tumor that reached from the oropharynx to the cervical junction. There was bone damage to the clivus as well as the cervical spine at C1, requiring a spinal fusion. Thus, the tumor itself exhibited migratory and destructive characteristics in the patient. From the cultured cell line we characterized the ARF-8 cellular and EV proteomes, as well as the impacts of ARF-8 EVs on proteomes and secretomes of recipient cells (both ARF-8 and human osteoblasts) in autocrine and paracrine settings. Our proteomic analyses suggested roles for TGFB and cell-matrix interactions in EMT, and cell/extracellular matrix interactions in cell migration, consistent with a migratory or even a metastatic tumor phenotype. Our results demonstrated that ARF-8 tumor cell migration was dependent on general (RGD-based) integrin activity, and ARF-8 EVs could promote such migration. ARF-8 EVs also promoted proteomic/secretomic changes in human osteoblast cells, again with indications that cell-cell and cell-extracellular matrix interactions would be activated.

ARF-8 as a cell line grew slowly, with a peak doubling time of almost 3 weeks. It expresses the embryonic transcription factor brachyury/TBXT (seen in Western blots, **Fig 2**), which is critical in vertebrate mesoderm and notochord development^48^, and is considered a reliable marker of most chordomas^49,50^. ARF-8 cells also express 10 different keratins, as well as 6 different S100A proteins, along with high quantities of vimentin (**Supp Table 2**), all classes of proteins that are also associated with chordoma diagnosis^2,51^. The Chordoma Foundation as a repository lists 26 patient-derived chordoma cell lines, emphasizing the rare nature of these cancers and the limited resources available to researchershttps://www.chordomafoundation.org/researchers/disease-models/

Extracellular vesicles (EVs) and other cell-released nanoparticles are increasingly recognized as critical communicators in biology in general, and in cancers in particular^52^. Despite the rapidly-expanding literature on the subject, as of this writing, there is only one publication on chordoma EVs/exosomes^21^. The authors isolated EVs, which they called “exosomes”, from conditioned media of the MUG-Chor1 cell line^53^ by a precipitation method, which yielded several larger-diameter (200 to > 400nm) populations of particles based on NTA; this was also the totality of the EV characterization. Here, we used more extensive means to isolate vesicles and conducted additional characterizations (NTA, TEM, ExoCheck, ExoView) along with what may be the first proteomic analysis of chordoma EVs (**Fig 2**, **3; Supp Table 1**). Over half of these EV proteins are classified as derived from the plasma membrane and extracellular space (**Fig 2**), and it is notable that the EVs contain 14 collagen proteins, 4 thrombospondins, 4 laminins, fibronectin, tenascins C and X, elastin, and heparan sulfate and chondroitin sulfate proteoglycans (HSPG2 and CSPG4). Other proteoglycans including aggrecan (ACAN), decorin (DCN), biglycan (BGN), versican (VCAN), osteomodulin (OMD), agrin (AGRN), SPARC/osteonectin, CWCV and Kazal-like domains proteoglycan 1 (SPOCK1), lumican (LUM), and glypican-1 (GPC1) are also present (**Supp Table 1**). Thus, extracellular matrix components are heavily represented in ARF-8 EVs. Pathway analysis reveals cell-cell and cell-matrix functional proclivities, in particular involving integrins (**Fig 2**). We again mention the presence of 5 integrin subunits (ITGB1, ITGA11, ITGAV, ITGA2, and ITGBL1) found in the EVs, which also implies promotion of EMT in conjunction with TGFB in the EVs (**Fig 2**). We also note the detection of brachyury/TBXT in EVs/extracellular particles from ARF-8 cells (**Fig 2**). As EVs are considered potential “liquid biopsy” biomarker sources in blood^54^, one speculates that TBXT in patient blood EVs could be a means of disease monitoring.

In autocrine (ARF-8 cells incubated with ARF-8 EVs) and paracrine (hOB cells incubated with ARF-8 EVs) protocols, we followed the changes in proteomic and secretomic outcomes of cells untreated or EV-treated. Common upregulated pathways and networks following EV treatment included glypican, syndecan, and integrin mediated signaling, and enhancement of tumorigenicity for EV-treated ARF-8 cells, with tumor-like phenotypic pathways prevalent in EV-treated hOB cells. Cell movement, migration, and invasion of tumor cells were also common themes (**Figs 4,7**). For the ARF-8 EV-stimulated secretome (**Fig 5**), we found high-scoring cell movement canonical pathways, which culminated in demonstrating that ARF-8 EVs are migration promoters (**Fig 5**). ARF-8 cells are quite motile even under minimal attractant conditions compared to their slow growth rates (**Fig 5 F,G**). It is possible that proteases released by ARF-8 cells following EV treatment, or matrix metalloprotease/collagenase activities directly on ARF-8 EVs themselves (**Fig 5 B,E**), may enhance cell migration. We identified 14 different collagens from ARF-8 cells (**Supp Table 2**), suggesting that extracellular protease release, particularly of MMPs, following EV treatment may target those extracellular matrix components to increase cellular migratory capacity.

ARF-8 proteomic and secretomic analyses, along with ARF-8 EV proteomics, were notable for the cell-cell and cell-matrix pathways derived from those data. The preponderance of integrin-related features, migratory pathways and networks led us to test ARF-8 cell reliance on RGD-dependent cell migration, where we found that 1 μM RGD peptide dramatically impaired ARF-8 cell migration (**Fig 6**) but did not induce cell death (which was seen at 10 μM and 100 μM doses, not shown). Integrins have long been known as important players in nearly all aspects of cancer progression^55^, but only one report has shown relevance in chordomas^56^. There, the authors found that antibody blocking of ITGB1 could induce cell death in U-CH1 cells and was additive with radiation to increase cell death in “cohesive cluster” cultured phenotypes that were particularly resistant to radiation. There have been decades of intense pre-clinical study, refinement, and clinical attempts to target integrins in cancer and other disease areas^57,58^. While there remain many challenges, targeting integrins in chordoma, therapeutically or as an imaging modality, seems to be an unexplored avenue.

The presence of TGFB in in ARF-8 EVs (**Supp Table 1**; **Fig 6**) and its recurring themes throughout our analyses (**Figs 3,6**, **7**), along with the aforementioned integrin subplots, led us to ask if anticipated relationships between these entities^57,59,60^ could resonate in the process of EMT in ARF-8 cells. Vasaikar et al^34^ developed the “EMTome” as tool to seek EMT/MET signatures in datasets, which we mined with our proteomic changes in ARF-8 cells treated with their own EVs. IPA applications produced a Graphical Summary (**Fig 6**, summary statement **Supp Fig 4**) that again placed TGFB1 at a major central node with associated SMAD and STAT signaling pathways activated. As the ARF-8 tumor had expanded extensively in the clivus and cervical spine, it was likely to have both EMT and MET expression components since the tumor presumably had reached its new niche. Curiously, the Graphical Summary also depicted Bone Mineralization and Osteoblast Differentiation networks as activated, which might result from the bony niche in which the resected tumor resided.

That information led us to examine osteoblasts as surrogates for the chordoma TME relative to the tumor’s clival niche. Lopez-Cuevas et al^35^ developed a zebrafish chordoma model by RAS-transforming notochord cells that led to destabilized spinal columns with activated osteoclasts, inflammatory processes, reduced bone density, and disorganized osteoblasts and collagen layers. This suggests that osteoblasts may be influenced by chordoma-like cells. These in vivo effects may be reflected in our proteomic analyses of cells treated with chordoma EVs, both the ARF-8 cells and the hOB cells with the extracellular and cell membrane-mediated signaling events, the remodeling pathway implications, and the overall cancerous environments suggested. In an analogous fashion (and noted above), Aydemir et al^21^ treated nucleus pulposus (NP) cells, long thought to be the cells of origin of chordomas^61^, with EVs (described as “exosomes”, but largely uncharacterized) from the MUG-Chor1 chordoma cell line. The exosomes promoted some phenotypic changes in the NP cells, as well as a mild increase in migration capacity. After a prolonged wait (140 days) after treating the NP cells, those cells showed some mRNA changes in stem cell markers, apoptotic markers, migration/invasion markers, and some tumor markers. From both that previous work and our work here, the implications are that chordoma EVs may induce microenvironment cells to take on tumor characteristics, as we have shown previously for glioblastoma EVs and astrocytes as cells of the TME^22^.

As chordomas are consistently referred to as chemo-resistant, novel molecular targets are sought^47,62^, which include the platelet-derived growth factor receptors (PDGFRs) targeted by imatinib and dasatinib; epidermal growth factor receptor (EGFR), targeted by erlotinib, lapatinib, gefitinib, and cetuximab; and anti-angiogenics such as sorafenib, pazopanib, and sunitinib to target vascular endothelial growth factor receptor (VEGFR). ARF-8 cells express PDGFRA and B (the EVs express PDGFRA) (**Supp Tables 1,2**), and the cells secrete PDGFA upon EV stimulation (**Fig 5**). Curiously, hOB cells also secrete PDGFA under both untreated and EV-stimulated conditions (**Fig 8**). ARF-8 cells also express EGFR (**Supp Table 2**), and both ARF-8 and hOBs (particularly ARF-8) secrete EGF further upon ARF-8 EV stimulation (**Figs 5,8**). VEGFR was not detected in our proteomic studies, but both ARF-8 and hOBs constitutively secrete VEGF. Thus, ARF-8 cells and their EVs could provide an interesting therapeutic model in the context of osteoblasts to test known or novel chemotherapies.

The bone-impacting nature of chordomas warrants attention. Bone destruction has long been associated with chordomas, along with elements of re-calcification seen surgically^63^ but also in imaging in spinal and clival settings^64,65^ up to this day^66^. Recently, a study revisited the chordoma bone-destructive invasive fronts to find cells expressing brachyury, tartrate-resistant acid phosphatase (TRAP/ACP5), and cathepsin K (CatK/CTSK), similar to osteoclasts in bone resorption capacities^67^. Cell line studies using the JHC7 chordoma line recapitulated features of activated osteoclasts in response to RANKL/TNFSF11, an osteoclastic cytokine. Matrix remodeling enzymes involved in osteogenesis are indeed expressed by chordomas^68^, and ARF-8 cells express a variety of such enzymes, including cathepsins A, B, D, K, L, and X/P/Z (from proteomics, **Supp Table 2**) along with secreted cathepsins A, B, C, D, E, L, S, V, and X/Z/P (from arrays, **Fig 5**). Further, MMP2 is highly expressed and secreted (**Fig 5**, but also on EVs, **Supp Table 1**), as well as the PLAU/PLAUR combination (**Fig 5**). Thus, ARF-8 cells and their EVs appear primed to alter the extracellular matrix and possibly may be involved in bone remodeling.

The prevalence of TGFB and integrins in these pathways suggests these molecules as potential therapeutic targets. Furthermore, ARF-8 EVs’ effects on both cancer cells and osteoblasts underscore the importance of EV-mediated communication in shaping the tumor microenvironment, particularly in bone-destructive chordomas. Our findings advocate for further exploration into the roles of EVs, integrins, and TGFB as critical players in chordoma progression, with implications for potential therapeutic interventions.

## CONCLUSION

We present a new chordoma cell line, ARF-8, derived from an extensive clival chordoma that invasively spread to the cervical spine. We have characterized the cells and the exosomes/extracellular vesicles from those cells, as well as the impacts the EVs had on the ARF-8 cells themselves along with effects on human osteoblasts as surrogate cells of the bony tumor microenvironment. This work presents novel insights into chordoma biology, particularly the role of chordoma EVs with their impacts in promoting tumorigenic behaviors in autocrine and paracrine settings. We note that the EVs possess strong extracellular matrix qualities and the abilities to promote ECM and cell-cell interactions, leading to EMT in the recipient cancer cells. Further, ARF-8 cell migration is integrin-mediated, and was promoted by ARF-8 EVs. In sum, our research with this novel cell line suggests a more critical look the roles of EVs in chordoma biology in general, with a focus on integrins and TGFB as possible targets for chordoma therapeutics in particular. This research represents a step forward in understanding the intricate dynamics between chordoma cells and their microenvironment, opening avenues for new therapeutic approaches targeting EV-mediated pathways.

---

Supplemental Figure 2. Ingenuity Pathway Analysis Summary for ARF-8 EV proteomics

Supplemental Figure 3. Ingenuity Pathway Analysis Summary for ARF-8 cell proteomics +/- ARF- 8 EVs

Supplemental Figure 4. Ingenuity Pathway Analysis Summary for ARF-8 EMT proteomics

Supplemental Figure 5. Ingenuity Pathway Analysis Summary for hOB cell proteomics +/- ARF-8 EVs

**Supplemental Figure 1.**
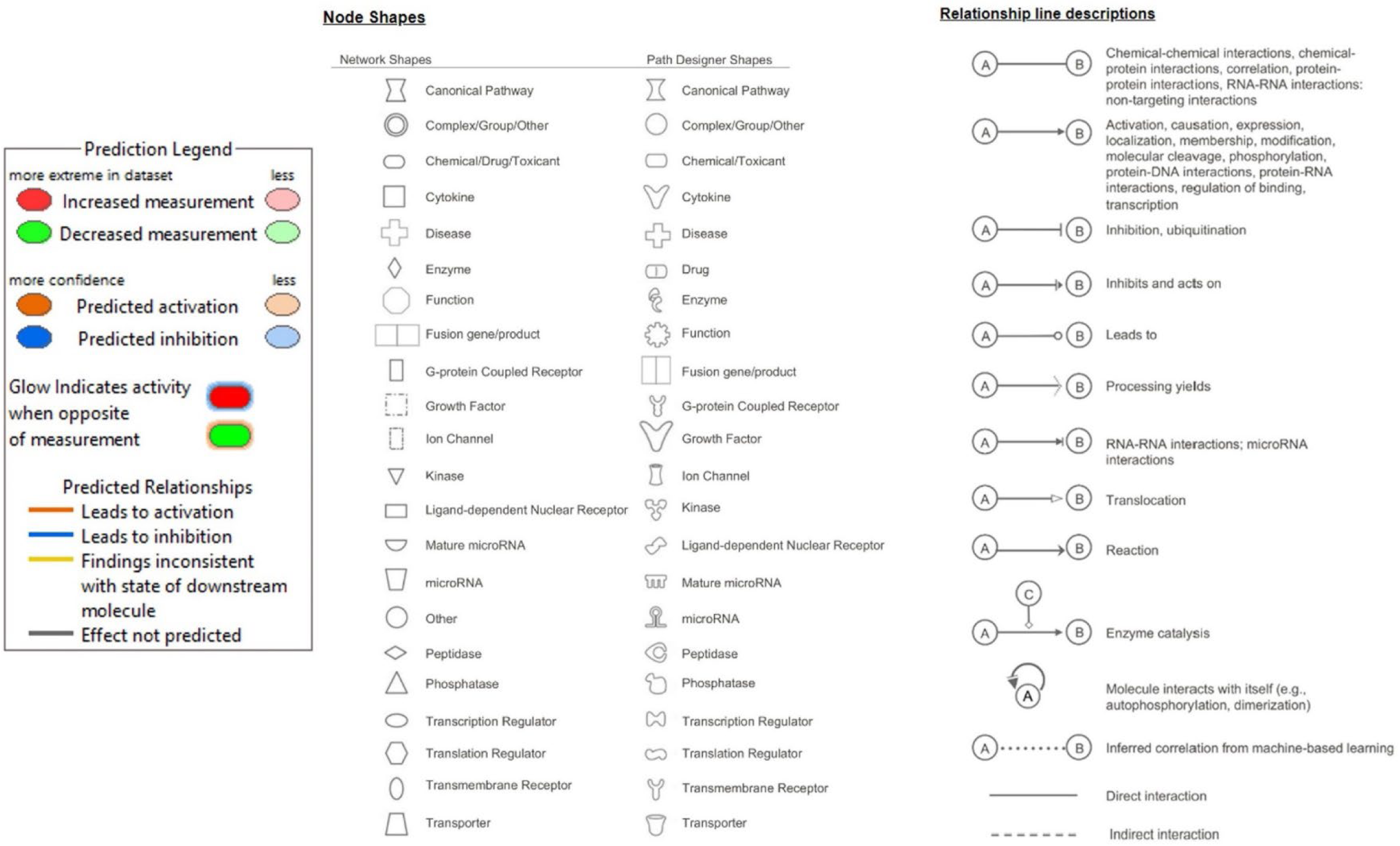
Ingenuity Pathway Analysis (IPA) network legends, shapes, and edge descriptions.

