## Supplementary material for "Extracellular vesicles from a novel chordoma cell line, ARF-8, promote tumorigenic microenvironmental changes when incubated with the parental cells and with human osteoblasts": Supp Fig 2

INGENUITY
PATHWAY ANALYSIS

Analysis Name: ARF-8 XO proteins for IPA - 2021-07-29 01:13 PM
Analysis Creation Date: 2021-07-29
Build version: exported
Content version: 65367011 (Release Date: 2021-06-04)

### Experiment Metadata

| Name | Value |
|---|---|

### Analysis Settings

Reference set: Ingenuity Knowledge Base (Genes + Endogenous Chemicals)
Relationship to include: Direct and Indirect
Includes Endogenous Chemicals
Optional Analyses: My Pathways My List

Filter Summary:
Consider only molecules and/or relationships where
(species = Uncategorized OR Human OR Rat OR Mouse) AND
(confidence = Experimentally Observed) AND
(tissues/cell lines = Stomach OR Thalamus OR U937 OR Activated CD56dim NK cells OR HeLa OR Ovary OR Neurons not otherwise specified OR Other Bone marrow cells OR M14 OR Peripheral blood lymphocytes OR Other Tissues and Primary Cells OR HL-60 OR Other Neurons OR COLO205 OR UO-31 OR Mesenchymal stem cells OR Smooth muscle cells not otherwise specified OR Immune cell lines not otherwise

(c) 2000-2024 QIAGEN. All rights reserved.

specified OR Dendritic cells not otherwise specified OR Activated Vd1 Gamma-delta T cells OR Intraepithelial T lymphocytes OR Hep3B OR Other Immune cell lines OR Effector T cells OR Other Macrophage Cancer Cell Lines OR Prostate Cancer Cell Lines not otherwise specified OR Cervical cancer cell line not otherwise specified OR Melanoma Cell Lines not otherwise specified OR Amygdala OR Peripheral blood monocytes OR Kidney cell lines not otherwise specified OR Cells not otherwise specified OR Spleen OR Subventricular Zone OR Mature monocyte-derived dendritic cells OR Cardiomyocytes OR H460 OR Osteosarcoma Cell Lines not otherwise specified OR T47-D OR Mononuclear leukocytes not otherwise specified OR Other Nervous System OR SK-OV-3 OR Plasmacytoid dendritic cells OR Retina OR Purkinje cells OR SR OR Activated CD56bright NK cells OR Endothelial cells not otherwise specified OR Vd1 Gamma-delta T cells OR P19 OR Macrophage Cancer Cell Lines not otherwise specified OR Striatum OR Langerhans cells OR Megakaryocytes OR Stem cells not otherwise specified OR RXF-393 OR SW-480 OR Prostate Gland OR Other Smooth muscle cells OR PANC-1 OR U87MG OR Eosinophils OR Stromal cells OR Other Epithelial cells OR LNCaP cells OR SK-MEL-28 OR BT-474 OR Neutrophils OR Breast Cancer Cell Lines not otherwise specified OR INS-1 OR Microvascular endothelial cells OR KM-12 OR Other B lymphocytes OR HCT-15 OR Effector memory helper T cells OR Memory T lymphocytes not otherwise specified OR A2780 OR SNB-75 OR HMC-1 OR Other Cervical cancer cell line OR Esophagus OR Other Lymphocytes OR Keratinocytes OR Central memory helper T cells OR Peritoneal macrophages OR Pre-B lymphocytes OR Other Prostate Cancer Cell Lines OR Other Pancreatic Cancer Cell Lines OR Oocytes OR Monocytes not otherwise specified OR Trigeminal Ganglion OR Min6 OR SW-620 OR PBMCs OR RKO OR Neuroblastoma Cell Lines not otherwise specified OR Thyroid Gland OR OVCAR-3 OR HOP-62 OR OVCAR-5 OR BDCA-1+ dendritic cells OR Natural T-regulatory cells OR Placenta OR 3T3-L1 cells OR Substantia Nigra OR 786-0 OR Dermis OR Memory B cells OR Other Ovarian Cancer Cell Lines OR CD34+ cells OR Cornea OR Calvaria OR Pyramidal neurons OR Lymphoma Cell Lines not otherwise specified OR SK-MEL-5 OR Granulocytes not otherwise specified OR Other Kidney cell lines OR Murine NKT cells OR Naive B cells OR Fibroblast cell lines not otherwise specified OR Other Macrophages OR Myeloid dendritic cells OR Other Dendritic cells OR Plasma cells OR Pheochromocytoma cell lines not otherwise specified OR Cell Line not otherwise specified OR Forestomach OR Osteoblasts OR Other Memory T lymphocytes OR Effector memory RA+ cytotoxic T cells OR Nucleus Accumbens OR Cerebral Cortex OR Fibroblasts OR MDA-MB-361 OR Other Stem cells OR HepG2 OR UACC-62 OR UACC-257 OR MDA-MB-231 OR PC-3 OR Caudate Nucleus OR Cerebral Ventricles OR Hepatoma Cell Lines not otherwise specified OR Dorsal Root Ganglion OR Epidermis OR Cos-7 cells OR NCI-H332M OR Other Kidney Cancer Cell Lines OR U2OS OR Thymus OR CD56dim NK cells OR Other Mononuclear leukocytes OR Myeloma Cell Lines not otherwise specified OR Mast cells OR Microglia OR Other Monocytes OR Pancreas OR CCRF-CEM OR Activated helper T cells OR Colon Cancer Cell Lines not otherwise specified OR NT2/D1 OR Corpus Callosum OR MCF7 OR Blood platelets OR BA/F3 OR Pituitary Gland OR Other Organ Systems OR PC-12 cells OR Large Intestine OR Vascular smooth muscle cells OR Effector memory cytotoxic T cells OR Bone marrow-derived macrophages OR Th1 cells OR Other Immune cells OR BT-549 OR RPMI-8266 OR Skin OR Th17 cells OR SK-MEL-2 OR A375 OR Adipocytes OR B lymphocytes not otherwise specified OR NB4 OR Other NK cells OR Bone marrow-derived dendritic cells OR LOX IMVI OR Putamen OR Adipose OR Hippocampus OR Other Cell Line

(c) 2000-2024 QIAGEN. All rights reserved.

OR Ventricular Zone OR SN12C OR Monocyte-derived macrophage OR Spinal Cord OR Immature monocyte-derived dendritic cells OR A549-ATCC OR Immune cells not otherwise specified OR Hepatocytes OR Organ Systems not otherwise specified OR Other Granulocytes OR Brain OR Smooth Muscle OR RBL-2H3 OR Heart OR Bone marrow cells not otherwise specified OR MG-63 OR Nervous System not otherwise specified OR J-774A.1 OR Embryonic stem cells OR SF-539 OR Adrenal Gland OR MDA-MB-468 OR Cortical neurons OR Pro-B lymphocytes OR Other Lymphoma Cell Lines OR HUVEC cells OR Gray Matter OR Kidney OR MDA-N OR MEF cells OR Splenocytes OR NCI-H23 OR Lymphocytes not otherwise specified OR Cerebellum OR K-562 OR Sertoli cells OR SK-N-SH OR Kidney Cancer Cell Lines not otherwise specified OR EKVX OR Epithelial cells not otherwise specified OR MOLT-4 OR 293 cells OR Olfactory Bulb OR Hypothalamus OR Other Fibroblast cell lines OR Lung OR Other Melanoma Cell Lines OR Ovarian Cancer Cell Lines not otherwise specified OR Other Leukemia Cell Lines OR Other Teratocarcinoma Cell Lines OR IGROV1 OR TK-10 OR Testis OR Caco2 cells OR Thymocytes OR Trachea OR Lung Cancer Cell Lines not otherwise specified OR Other Endothelial cells OR Cartilage Tissue OR Lymph node OR Other Peripheral blood leukocytes OR HEL OR Other T lymphocytes OR Crypt OR CD4+ T-lymphocytes OR Cytotoxic T cells OR Jurkat OR Parietal Lobe OR NCI-ADR-RES OR Teratocarcinoma Cell Lines not otherwise specified OR Skeletal Muscle OR White Matter OR Other Colon Cancer Cell Lines OR NK cells not otherwise specified OR HS 578T OR Medulla Oblongata OR Other Monocyte-derived dendritic cells OR SF-295 OR Sciatic Nerve OR Activated Vd2 Gamma-delta T cells OR A498 OR HCT-116 OR Other Osteosarcoma Cell Lines OR RAW 264.7 OR Th2 cells OR Pancreatic Cancer Cell Lines not otherwise specified OR Vd2 Gamma-delta T cells OR Hematopoietic progenitor cells OR ACHN OR Swiss 3T3 cells OR NCI-H522 OR U266 OR Other CNS Cell Lines OR Other Lung Cancer Cell Lines OR Granule cells OR Mammary Gland OR Liver OR Melanocytes OR Other Cells OR Other Pheochromocytoma cell lines OR Beta islet cells OR Monocyte-derived dendritic cells not otherwise specified OR Lens OR Bladder OR NIH/3T3 cells OR J774 OR Other Neuroblastoma Cell Lines OR MDA-MB-435 OR MALME-3M OR Uterus OR Central memory cytotoxic T cells OR HCC-2998 OR NCI-H226 OR Granulosa cells OR HOP-92 OR Macrophages not otherwise specified OR Chondrocytes OR Other Hepatoma Cell Lines OR SF-268 OR CD56bright NK cells OR HT29 OR Other Myeloma Cell Lines OR Brainstem OR CAKI-1 OR Choroid Plexus OR Granule Cell Layer OR Tissues and Primary Cells not otherwise specified OR Other Breast Cancer Cell Lines OR OVCAR-8 OR T lymphocytes not otherwise specified OR BDCA-3+ dendritic cells OR Small Intestine OR U251 OR Peripheral blood leukocytes not otherwise specified OR HuH7 OR WEHI-231 OR Astrocytes OR CNS Cell Lines not otherwise specified OR DU-145 OR THP-1 OR Leukemia Cell Lines not otherwise specified OR OVCAR-4 OR Salivary Gland OR Naive helper T cells) AND

(mol. types = biologic drug OR canonical pathway OR chemical - endogenous mammalian OR chemical - endogenous non-mammalian OR chemical - kinase inhibitor OR chemical - other OR chemical - protease inhibitor OR chemical drug OR chemical reagent OR chemical toxicant OR complex OR cytokine OR disease OR enzyme OR function OR fusion gene/product OR G-protein coupled receptor OR group OR growth factor OR ion channel OR kinase OR ligand-dependent nuclear receptor OR mature microRNA OR microRNA OR other OR peptidase OR phosphatase OR transcription regulator OR translation regulator OR transmembrane receptor OR transporter) AND

(c) 2000-2024 QIAGEN. All rights reserved.

(data sources = An Open Access Database of Genome-wide Association Results OR BIND OR BioGRID OR Catalogue Of Somatic Mutations In Cancer (COSMIC) OR Chemical Carcinogenesis Research Information System (CCRIS) OR Clinical Genome Resource (ClinGen) OR ClinicalTrials.gov OR ClinVar OR Cognia OR DIP OR DrugBank OR Gene Ontology (GO) OR GVK Biosciences OR Hazardous Substances Data Bank (HSDB) OR HumanCyc OR Ingenuity Expert Findings OR Ingenuity ExpertAssist Findings OR IntAct OR Interactome studies OR MIPS OR miRBase OR miRecords OR Mouse Genome Database (MGD) OR Obesity Gene Map Database OR Online Mendelian Inheritance in Man (OMIM) OR TarBase OR TargetScan Human)

### Top Canonical Pathways

| Name | p-value | Overlap |
|---|---|---|
| **GP6 Signaling Pathway** | 1.57E-20 | 17.4 % 23/132 |
| **Hepatic Fibrosis / Hepatic Stellate Cell Activation** | 1.10E-18 | 12.6 % 25/198 |
| **Actin Cytoskeleton Signaling** | 2.50E-13 | 8.8 % 22/249 |
| **Axonal Guidance Signaling** | 3.15E-12 | 5.7 % 29/512 |
| **Integrin Signaling** | 8.84E-11 | 8.4 % 18/214 |

### Top Upstream Regulators

#### Upstream Regulators

| Name | p-value | Predicted Activation |
|---|---|---|
| **TGFB1** | 6.14E-56 | |
| **beta-estradiol** | 1.46E-48 | |
| **AGT** | 2.80E-39 | |

(c) 2000-2024 QIAGEN. All rights reserved.

INGENUITY PATHWAY ANALYSIS

| | |
|---|---|
| **dexamethasone** | 1.79E-37 |
| **TP53** | 1.85E-36 |

#### Causal Network

| Name | p-value | Predicted Activation |
|---|---|---|
| **aprotinin** | 1.20E-62 | |
| **Thrombospondin** | 1.39E-55 | |
| **ferric chloride** | 1.06E-53 | |
| **sulpiride** | 9.09E-53 | |
| **MYCN** | 4.02E-52 | |

### Top Diseases and Bio Functions

#### Diseases and Disorders

| Name | p-value range | # Molecules |
|---|---|---|
| **Cancer** | 3.14E-12 - 2.47E-39 | 295 |
| **Organismal Injury and Abnormalities** | 3.14E-12 - 2.47E-39 | 297 |
| **Inflammatory Response** | 2.15E-12 - 5.80E-39 | 178 |
| **Reproductive System Disease** | 3.14E-12 - 9.65E-39 | 252 |
| **Gastrointestinal Disease** | 2.81E-12 - 1.16E-34 | 284 |

#### Molecular and Cellular Functions

(c) 2000-2024 QIAGEN. All rights reserved.

INGENUITY PATHWAY ANALYSIS

| Name | p-value range | # Molecules |
| --- | --- | --- |
| **Cellular Movement** | 2.09E-12 - 3.35E-57 | 179 |
| **Cellular Compromise** | 1.71E-22 - 5.80E-39 | 70 |
| **Protein Synthesis** | 1.51E-33 - 4.78E-38 | 94 |
| **Cell Death and Survival** | 2.66E-12 - 2.86E-35 | 177 |
| **Cell-To-Cell Signaling and Interaction** | 8.36E-13 - 4.51E-28 | 140 |

#### Physiological System Development and Function

| Name | p-value range | # Molecules |
| --- | --- | --- |
| **Tissue Development** | 2.22E-12 - 3.64E-58 | 168 |
| **Cardiovascular System Development and Function** | 2.37E-12 - 1.22E-49 | 140 |
| **Organismal Development** | 2.37E-12 - 9.02E-46 | 173 |
| **Hematological System Development and Function** | 1.32E-12 - 5.05E-27 | 93 |
| **Immune Cell Trafficking** | 1.32E-12 - 5.05E-27 | 83 |

### Top Tox Functions

#### Assays: Clinical Chemistry and Hematology

| Name | p-value range | # Molecules |
| --- | --- | --- |
| **Decreased Levels of Albumin** | 1.52E-01 - 1.21E-03 | 4 |
| **Increased Levels of Blood Urea Nitrogen** | 6.71E-03 - 6.71E-03 | 3 |

(c) 2000-2024 QIAGEN. All rights reserved.

INGENUITY PATHWAY ANALYSIS

| | | |
|---|---|---|
| **Increased Levels of Albumin** | 1.17E-02 - 1.17E-02 | 1 |
| **Increased Levels of Potassium** | 2.05E-02 - 2.05E-02 | 2 |
| **Increased Levels of Bilirubin** | 2.33E-02 - 2.33E-02 | 1 |

### Cardiotoxicity

| Name | p-value range | # Molecules |
|---|---|---|
| **Cardiac Enlargement** | 3.38E-01 - 2.39E-11 | 38 |
| **Cardiac Arrythmia** | 2.47E-01 - 1.77E-08 | 21 |
| **Cardiac Dilation** | 2.73E-01 - 5.08E-08 | 23 |
| **Cardiac Infarction** | 1.62E-01 - 1.99E-07 | 18 |
| **Cardiac Fibrosis** | 1.11E-01 - 2.31E-07 | 17 |

### Hepatotoxicity

| Name | p-value range | # Molecules |
|---|---|---|
| **Liver Hyperplasia/Hyperproliferation** | 2.55E-01 - 7.30E-21 | 176 |
| **Hepatocellular carcinoma** | 1.91E-01 - 9.54E-16 | 71 |
| **Liver Fibrosis** | 1.61E-01 - 6.88E-09 | 25 |
| **Liver Cirrhosis** | 1.61E-01 - 3.07E-08 | 20 |
| **Liver Proliferation** | 2.47E-01 - 1.46E-06 | 12 |

### Nephrotoxicity

(c) 2000-2024 QIAGEN. All rights reserved.

INGENUITY PATHWAY ANALYSIS

| Name | p-value range | # Molecules |
|---|---|---|
| **Glomerular Injury** | 1.42E-01 - 1.35E-11 | 31 |
| **Renal Damage** | 2.47E-01 - 1.10E-09 | 25 |
| **Renal Tubule Injury** | 2.47E-01 - 1.10E-09 | 16 |
| **Nephrosis** | 1.42E-01 - 3.54E-07 | 13 |
| **Renal Necrosis/Cell Death** | 1.72E-01 - 6.31E-07 | 24 |

### Top Regulator Effect Networks

### Top Networks

| ID | Associated Network Functions | Score |
|---|---|---|
| **1** | Connective Tissue Disorders, Dermatological Diseases and Conditions, Developmental Disorder | 39 |
| **2** | Tissue Development, Cardiovascular System Development and Function, Cancer | 36 |
| **3** | Cellular Assembly and Organization, Cellular Function and Maintenance, Tissue Development | 36 |
| **4** | Tissue Development, Cancer, Connective Tissue Disorders | 34 |

(c) 2000-2024 QIAGEN. All rights reserved.

INGENUITY PATHWAY ANALYSIS

| | | | |
|---|---|---|---|
| 5 | | Cell Morphology, Embryonic Development, Hair and Skin Development and Function | 34 |

### Top Tox Lists

| Name | p-value | Overlap |
|---|---|---|
| Hepatic Fibrosis | 2.75E-22 | 10.0 % 35/351 |
| Genes associated with Chronic Allograft Nephropathy (Human) | 5.80E-11 | 38.1 % 8/21 |
| Acute Renal Failure Panel (Rat) | 1.43E-10 | 17.7 % 11/62 |
| Renal Necrosis/Cell Death | 1.13E-07 | 4.0 % 25/625 |
| Cardiac Fibrosis | 4.72E-07 | 5.4 % 16/295 |

### Top My Lists

### Top My Pathways

### Top ML Disease Pathways

### Top Analysis-Ready Molecules

(c) 2000-2024 QIAGEN. All rights reserved.
