## Supplementary material for "Extracellular vesicles from a novel chordoma cell line, ARF-8, promote tumorigenic microenvironmental changes when incubated with the parental cells and with human osteoblasts": Supp Fig 3

INGENUITY
PATHWAY ANALYSIS

Analysis Name: ARF8 XOtx v Cont 03Sept for IPA - 2021-09-03 03:26 PM
Analysis Creation Date: 2021-09-03
Build version: exported
Content version: 65367011 (Release Date: 2021-06-04)

### Experiment Metadata

### Top Canonical Pathways

| Name | p-value | Overlap |
|---|---|---|
| **EIF2 Signaling** | 2.19E-07 | 7.5 % 17/227 |
| **Tumor Microenvironment Pathway** | 5.05E-07 | 8.0 % 15/188 |
| **Glucocorticoid Receptor Signaling** | 1.75E-06 | 4.6 % 27/591 |
| **CSDE1 Signaling Pathway** | 3.32E-06 | 14.3 % 8/56 |
| **Caveolar-mediated Endocytosis Signaling** | 4.49E-06 | 11.7 % 9/77 |

### Top Upstream Regulators

#### Upstream Regulators

| Name | p-value | Predicted Activation |
|---|---|---|
| **beta-estradiol** | 3.48E-15 | |
| **TP53** | 2.76E-14 | |
| **ESR1** | 1.17E-13 | |
| **OSM** | 3.88E-12 | |

(c) 2000-2024 QIAGEN. All rights reserved.

| | |
|---|---|
| **CST5** | 1.38E-11 |

#### Causal Network

| Name | p-value | Predicted Activation |
|---|---|---|
| **Rasgrp** | 2.75E-21 | |
| **DNAJB1** | 1.61E-20 | |
| **SP4** | 4.50E-20 | Activated |
| **SYVN1** | 4.61E-20 | |
| **SVIL** | 6.32E-20 | |

### Top Diseases and Bio Functions

#### Diseases and Disorders

| Name | p-value range | # Molecules |
|---|---|---|
| **Cancer** | 2.27E-05 - 1.66E-37 | 404 |
| **Organismal Injury and Abnormalities** | 2.78E-05 - 1.66E-37 | 408 |
| **Endocrine System Disorders** | 1.17E-05 - 5.72E-33 | 357 |
| **Gastrointestinal Disease** | 2.73E-05 - 3.29E-25 | 365 |
| **Neurological Disease** | 2.74E-05 - 4.59E-21 | 310 |

#### Molecular and Cellular Functions

(c) 2000-2024 QIAGEN. All rights reserved.

| Name | p-value range | # Molecules |
|---|---|---|
| Cellular Assembly and Organization | 1.25E-05 - 3.79E-17 | 147 |
| Cellular Function and Maintenance | 2.51E-05 - 3.79E-17 | 179 |
| Cell Death and Survival | 2.83E-05 - 6.74E-15 | 188 |
| Cellular Movement | 1.73E-05 - 1.22E-13 | 141 |
| Cell Morphology | 2.75E-05 - 2.90E-12 | 138 |

#### Physiological System Development and Function

| Name | p-value range | # Molecules |
|---|---|---|
| Organismal Survival | 2.31E-06 - 1.16E-16 | 148 |
| Organismal Development | 2.78E-05 - 5.07E-12 | 181 |
| Cardiovascular System Development and Function | 2.78E-05 - 4.18E-11 | 94 |
| Tissue Development | 1.96E-05 - 4.69E-11 | 156 |
| Nervous System Development and Function | 1.33E-05 - 6.20E-10 | 105 |

### Top Tox Functions

#### Assays: Clinical Chemistry and Hematology

| Name | p-value range | # Molecules |
|---|---|---|
| Increased Levels of Albumin | 1.79E-01 - 4.76E-03 | 4 |
| Increased Levels of Red Blood Cells | 1.11E-01 - 1.11E-01 | 4 |

(c) 2000-2024 QIAGEN. All rights reserved.

INGENUITY PATHWAY ANALYSIS

| | | |
|---|---|---|
| Increased Levels of LDH | 1.55E-01 - 1.55E-01 | 2 |
| Decreased Levels of Hematocrit | 1.65E-01 - 1.65E-01 | 1 |
| Increased Levels of Hematocrit | 2.23E-01 - 2.23E-01 | 3 |

#### Cardiotoxicity

| Name | p-value range | # Molecules |
|---|---|---|
| Cardiac Dilation | 7.11E-02 - 1.09E-06 | 25 |
| Cardiac Enlargement | 1.92E-01 - 1.09E-06 | 35 |
| Cardiac Dysfunction | 5.82E-01 - 4.02E-06 | 16 |
| Cardiac Necrosis/Cell Death | 9.38E-02 - 2.20E-04 | 16 |
| Cardiac Fibrosis | 2.68E-01 - 2.35E-04 | 15 |

#### Hepatotoxicity

| Name | p-value range | # Molecules |
|---|---|---|
| Liver Hyperplasia/Hyperproliferation | 4.73E-01 - 2.88E-11 | 200 |
| Hepatocellular carcinoma | 4.73E-01 - 1.11E-06 | 65 |
| Liver Necrosis/Cell Death | 1.44E-01 - 6.57E-04 | 16 |
| Liver Steatosis | 5.71E-01 - 1.19E-03 | 18 |
| Liver Fibrosis | 3.26E-01 - 2.94E-03 | 18 |

#### Nephrotoxicity

(c) 2000-2024 QIAGEN. All rights reserved.

INGENUITY PATHWAY ANALYSIS

| Name | p-value range | # Molecules |
|---|---|---|
| **Glomerular Injury** | 5.15E-01 - 4.82E-06 | 19 |
| **Renal Inflammation** | 1.00E00 - 1.55E-03 | 8 |
| **Renal Nephritis** | 1.00E00 - 1.55E-03 | 8 |
| **Renal Damage** | 1.00E00 - 2.53E-03 | 10 |
| **Renal Tubule Injury** | 7.88E-02 - 2.53E-03 | 8 |

### Top Regulator Effect Networks

| ID | Regulators | Disease & Functions | Consistency Score |
|---|---|---|---|
| **1** | CD5,GPER1,GSK3 (family),IL3,MAP3K14 (+3 more) | Apoptosis,Cell survival (+3 more) | 14.8 |
| **2** | GSK3 (family),MAP3K14,MLXIPL,P70 S6K (family),PKM | Apoptosis,Cell survival (+3 more) | 12.746 |
| **3** | FLI1,IKBKB,PKM | Homing of cells,Invasion of cells (+1 more) | 8.0 |
| **4** | ASXL1,IGF1R,mir-15 (includes others) (+3 more) | Cell movement of tumor cell lines (+7 more) | 7.903 |
| **5** | PKM | Cell proliferation of tumor cell lines (+3 more) | 7.757 |

### Top Networks

| ID | Associated Network Functions | Score |
|---|---|---|
| **1** | RNA Post-Transcriptional Modification, Cancer, Organismal Injury and Abnormalities | 55 |

(c) 2000-2024 QIAGEN. All rights reserved.

INGENUITY PATHWAY ANALYSIS

| | | |
|---|---|---|
| **2** | Hematological Disease, Nutritional Disease, Metabolic Disease | 50 |
| **3** | Protein Synthesis, Gene Expression, Cellular Assembly and Organization | 45 |
| **4** | Carbohydrate Metabolism, Nucleic Acid Metabolism, Small Molecule Biochemistry | 38 |
| **5** | Cancer, Dermatological Diseases and Conditions, Organismal Injury and Abnormalities | 35 |

### Top Tox Lists

| Name | p-value | Overlap |
|---|---|---|
| **Genes associated with Chronic Allograft Nephropathy (Human)** | 1.84E-05 | 23.8 % 5/21 |
| **Cardiac Necrosis/Cell Death** | 6.96E-05 | 5.1 % 16/316 |
| **Cardiac Fibrosis** | 1.11E-04 | 5.1 % 15/295 |
| **Liver Necrosis/Cell Death** | 1.19E-04 | 4.8 % 16/331 |
| **Hepatic Fibrosis** | 7.05E-04 | 4.3 % 15/351 |

### Top My Lists

(c) 2000-2024 QIAGEN. All rights reserved.

INGENUITY PATHWAY ANALYSIS

### Top My Pathways

### Top ML Disease Pathways

### Top Analysis-Ready Molecules

#### Expr Log Ratio

| Molecules | Expr. Value | Chart |
|---|---|---|
| AAK1 | ↑ 10.000 | |
| ABHD12 | ↑ 10.000 | |
| ADAMTS4 | ↑ 10.000 | |
| ADGRL2 | ↑ 10.000 | |
| AKAP11 | ↑ 10.000 | |
| AKT1S1 | ↑ 10.000 | |
| AKT2 | ↑ 10.000 | |
| ANKH | ↑ 10.000 | |
| ANP32A | ↑ 10.000 | |
| AP1S1 | ↑ 10.000 | |

#### Expr Log Ratio

(c) 2000-2024 QIAGEN. All rights reserved.

| Molecules | Expr. Value | Chart |
|---|---|---|
| **XRN2** | ↓ -10.000 | |
| **VTI1B** | ↓ -10.000 | |
| **UAP1L1** | ↓ -10.000 | |
| **TRMT6** | ↓ -10.000 | |
| **THOP1** | ↓ -10.000 | |
| **STK17B** | ↓ -10.000 | |
| **SRP9** | ↓ -10.000 | |
| **SORD** | ↓ -10.000 | |
| **SNAP29** | ↓ -10.000 | |
| **SF1** | ↓ -10.000 | |

(c) 2000-2024 QIAGEN. All rights reserved.
