## Supplementary material for "Extracellular vesicles from a novel chordoma cell line, ARF-8, promote tumorigenic microenvironmental changes when incubated with the parental cells and with human osteoblasts": Supp Fig 4

INGENUITY
PATHWAY ANALYSIS

Analysis Name: ARF8 EMT cells&ALL for IPA#2 - 2023-09-10 08:53 AM
Analysis Creation Date: 2023-09-10
Build version: exported
Content version: 94302991 (Release Date: 2023-05-27)

### Experiment Metadata

| Name | Value |
|---|---|

### Analysis Settings

### Top Canonical Pathways

| Name | p-value | Overlap |
|---|---|---|
| Pulmonary Fibrosis Idiopathic Signaling Pathway | 1.08E-45 | 17.2 % 56/326 |
| Hepatic Fibrosis / Hepatic Stellate Cell Activation | 5.29E-36 | 20.6 % 40/194 |
| Wound Healing Signaling Pathway | 4.34E-30 | 15.5 % 39/252 |
| Hepatic Fibrosis Signaling Pathway | 1.79E-24 | 9.9 % 42/423 |
| Tumor Microenvironment Pathway | 2.12E-24 | 16.8 % 30/179 |

(c) 2000-2024 QIAGEN. All rights reserved.

### Top Upstream Regulators

#### Upstream Regulators

| Name | p-value | Predicted Activation |
|---|---|---|
| TGFB1 | 1.37E-113 | Activated |
| HRAS | 2.47E-70 | |
| KRAS | 3.49E-70 | Inhibited |
| beta-estradiol | 2.00E-68 | Activated |
| AGT | 2.93E-67 | Activated |

#### Causal Network

| Name | p-value | Predicted Activation |
|---|---|---|
| LY2109761 | 2.33E-100 | Inhibited |
| KLK14 | 1.13E-99 | Activated |
| TRPS1 | 3.11E-98 | Inhibited |
| TGFB1 | 3.21E-97 | Activated |
| ARID1A | 4.33E-73 | |

### Top Diseases and Bio Functions

(c) 2000-2024 QIAGEN. All rights reserved.

INGENUITY PATHWAY ANALYSIS

### Diseases and Disorders

| Name | p-value range | # Molecules |
|---|---|---|
| Cancer | 6.46E-14 - 2.11E-70 | 318 |
| Organismal Injury and Abnormalities | 7.26E-14 - 2.11E-70 | 318 |
| Reproductive System Disease | 6.46E-14 - 2.33E-48 | 287 |
| Tumor Morphology | 3.75E-14 - 2.90E-48 | 125 |
| Respiratory Disease | 5.48E-15 - 5.49E-46 | 240 |

### Molecular and Cellular Functions

| Name | p-value range | # Molecules |
|---|---|---|
| Cellular Movement | 3.36E-14 - 8.62E-89 | 234 |
| Cellular Growth and Proliferation | 6.21E-14 - 1.13E-54 | 236 |
| Cellular Assembly and Organization | 5.13E-14 - 8.11E-52 | 209 |
| Cellular Function and Maintenance | 7.69E-14 - 8.11E-52 | 218 |
| Cell Death and Survival | 7.69E-14 - 1.30E-51 | 212 |

### Physiological System Development and Function

| Name | p-value range | # Molecules |
|---|---|---|
| Cardiovascular System Development and Function | 6.21E-14 - 2.44E-78 | 178 |
| Organismal Development | 6.21E-14 - 2.41E-75 | 251 |
| Connective Tissue Development and Function | 6.21E-14 - 1.13E-54 | 186 |
| Tissue Development | 6.21E-14 - 1.13E-54 | 247 |

(c) 2000-2024 QIAGEN. All rights reserved.

INGENUITY PATHWAY ANALYSIS

| | | |
|---|---|---|
| Organismal Survival | 9.35E-20 - 4.94E-53 | 193 |

### Top Tox Functions

#### Assays: Clinical Chemistry and Hematology

| Name | p-value range | # Molecules |
|---|---|---|
| Increased Levels of Alkaline Phosphatase | 1.32E-02 - 1.27E-10 | 13 |
| Decreased Levels of Albumin | 5.19E-02 - 9.09E-06 | 5 |
| Increased Levels of LDH | 1.64E-02 - 1.38E-04 | 5 |
| Increased Levels of Albumin | 1.48E-01 - 3.57E-04 | 4 |
| Increased Levels of Hematocrit | 4.45E-04 - 4.45E-04 | 7 |

#### Cardiotoxicity

| Name | p-value range | # Molecules |
|---|---|---|
| Cardiac Dysfunction | 2.24E-01 - 5.80E-34 | 62 |
| Cardiac Enlargement | 3.38E-01 - 9.83E-23 | 58 |
| Cardiac Fibrosis | 1.36E-01 - 2.03E-18 | 33 |
| Cardiac Arrythmia | 5.26E-01 - 7.05E-13 | 28 |
| Cardiac Dilation | 3.38E-01 - 1.53E-12 | 32 |

#### Hepatotoxicity

(c) 2000-2024 QIAGEN. All rights reserved.

INGENUITY PATHWAY ANALYSIS

| Name | p-value range | # Molecules |
|---|---|---|
| **Liver Fibrosis** | 1.01E-01 - 8.27E-21 | 50 |
| **Hepatocellular carcinoma** | 3.47E-01 - 9.20E-20 | 92 |
| **Liver Hyperplasia/Hyperproliferation** | 3.47E-01 - 9.20E-20 | 191 |
| **Liver Proliferation** | 3.02E-01 - 1.96E-15 | 26 |
| **Liver Necrosis/Cell Death** | 6.44E-02 - 9.59E-12 | 25 |

#### Nephrotoxicity

| Name | p-value range | # Molecules |
|---|---|---|
| **Renal Proliferation** | 1.01E-01 - 3.38E-13 | 26 |
| **Glomerular Injury** | 4.51E-01 - 4.73E-12 | 44 |
| **Renal Hydronephrosis** | 4.55E-11 - 4.55E-11 | 14 |
| **Renal Necrosis/Cell Death** | 1.59E-01 - 4.66E-11 | 36 |
| **Renal Damage** | 1.35E-01 - 2.44E-10 | 31 |

### Top Regulator Effect Networks

| ID | Regulators | Disease & Functions | Consistency Score |
|---|---|---|---|
| **1** | bleomycin | Migration of carcinoma cell lines | 3.474 |
| **2** | YAP1 | Cell movement of lung cancer cell lines | 3.464 |
| **3** | bleomycin | Epithelial-mesenchymal transition | 3.357 |
| **4** | HMG20A | Organismal death | 3.336 |
| **5** | aldosterone | Size of body | 3.328 |

(c) 2000-2024 QIAGEN. All rights reserved.

### Top Networks

| ID | Associated Network Functions | Score |
| --- | --- | --- |
| **1** | Organ Development, Reproductive System Development and Function, Organismal Development | 38 |
| **2** | Cell Morphology, Embryonic Development, Hair and Skin Development and Function | 31 |
| **3** | Cellular Movement, Skeletal and Muscular System Development and Function, Cancer | 31 |
| **4** | Dermatological Diseases and Conditions, Inflammatory Disease, Inflammatory Response | 31 |
| **5** | Organismal Injury and Abnormalities, Organismal Survival, Cancer | 29 |

### Top Tox Lists

(c) 2000-2024 QIAGEN. All rights reserved.

INGENUITY PATHWAY ANALYSIS

| Name | p-value | Overlap |
|---|---|---|
| **Hepatic Fibrosis** | 4.45E-38 | 14.7 % 51/347 |
| **Cardiac Fibrosis** | 9.37E-19 | 9.5 % 33/349 |
| **Genes associated with Chronic Allograft Nephropathy (Human)** | 6.17E-18 | 57.1 % 12/21 |
| **Cardiac Hypertrophy** | 1.20E-15 | 8.0 % 31/387 |
| **Liver Proliferation** | 1.04E-14 | 9.2 % 26/283 |

### Top My Lists

### Top My Pathways

### Top ML Disease Pathways

| Name | p-value | Overlap |
|---|---|---|
| **Connective tissue dysplasia** | 3.01E-12 | 23.1 % 12/52 |
| **Dysplasia of skeleton** | 2.08E-11 | 23.4 % 11/47 |
| **Advanced malignant tumor** | 8.51E-11 | 25.0 % 10/40 |
| **Advanced stage tumor** | 8.51E-11 | 25.0 % 10/40 |
| **Corneal neovascularization** | 8.80E-11 | 31.0 % 9/29 |

### Top Analysis-Ready Molecules

**Expr Log Ratio**

(c) 2000-2024 QIAGEN. All rights reserved.

| Molecules | Expr. Value | Chart |
|---|---|---|
| SULF1 | ↑ 3.059 | |
| ECM2 | ↑ 3.016 | |
| FGF2 | ↑ 3.016 | |
| DNAJB4 | ↑ 3.000 | |
| ANGPTL4 | ↑ 2.031 | |
| C1S | ↑ 2.000 | |
| COMP | ↑ 2.000 | |
| FST | ↑ 2.000 | |
| HACL1 | ↑ 2.000 | |
| LAMA2 | ↑ 2.000 | |

**Expr Log Ratio**

| Molecules | Expr. Value | Chart |
|---|---|---|
| BMI1 | ↓ -15.673 | |
| MBNL1 | ↓ -11.551 | |
| FZD2 | ↓ -11.551 | |
| FUCA1 | ↓ -2.710 | |
| COL5A2 | ↓ -2.419 | |
| MAPKAP1 | ↓ -0.794 | |
| PLAUR | ↓ -0.299 | |
| KLK6 | ↓ -0.250 | |
| TFPI2 | ↓ -0.224 | |
| SLIT2 | ↓ -0.201 | |

(c) 2000-2024 QIAGEN. All rights reserved.

INGENUITY PATHWAY ANALYSIS
