## Supplementary material for "Extracellular vesicles from a novel chordoma cell line, ARF-8, promote tumorigenic microenvironmental changes when incubated with the parental cells and with human osteoblasts": Supp Fig 5

INGENUITY
PATHWAY ANALYSIS

Analysis Name: hOB plus XO log2 for IPA#3 Aug 2022all - 2022-08-17 07:19 AM
Analysis Creation Date: 2022-08-17
Build version: exported
Content version: 76765844 (Release Date: 2022-06-01)

### Top Canonical Pathways

| Name | p-value | Overlap |
|---|---|---|
| **EIF2 Signaling** | 4.48E-66 | 63.9 % 147/230 |
| **Regulation of eIF4 and p70S6K Signaling** | 2.04E-42 | 57.8 % 107/185 |
| **Protein Ubiquitination Pathway** | 5.92E-33 | 44.2 % 123/278 |
| **Integrin Signaling** | 8.11E-32 | 48.4 % 103/213 |
| **mTOR Signaling** | 8.69E-32 | 47.7 % 105/220 |

### Top Upstream Regulators

#### Upstream Regulators

| Name | p-value | Predicted Activation |
|---|---|---|
| **TP53** | 1.60E-107 | Activated |
| **MYC** | 9.02E-88 | Activated |
| **HNF4A** | 3.39E-86 | |
| **torin1** | 1.53E-69 | Inhibited |

(c) 2000-2024 QIAGEN. All rights reserved.

INGENUITY PATHWAY ANALYSIS

| | | |
|---|---|---|
| **GABA** | 2.65E-61 | Inhibited |

#### Causal Network

| Name | p-value | Predicted Activation |
|---|---|---|
| **Ep300/Pcaf** | 6.63E-137 | Activated |
| **SGI 1776** | 5.22E-133 | Inhibited |
| **Jmy-p300** | 7.10E-132 | Activated |
| **cadmium** | 3.62E-131 | Activated |
| **stigmatellin** | 4.61E-130 | Inhibited |

### Top Diseases and Bio Functions

#### Diseases and Disorders

| Name | p-value range | # Molecules |
|---|---|---|
| **Cancer** | 6.79E-22 - 0.00E00 | 3630 |
| **Organismal Injury and Abnormalities** | 1.42E-21 - 0.00E00 | 3676 |
| **Endocrine System Disorders** | 2.28E-22 - 4.10E-260 | 3174 |
| **Gastrointestinal Disease** | 6.79E-22 - 1.03E-217 | 3254 |
| **Neurological Disease** | 9.80E-22 - 4.77E-114 | 2716 |

#### Molecular and Cellular Functions

(c) 2000-2024 QIAGEN. All rights reserved.

INGENUITY PATHWAY ANALYSIS

| Name | p-value range | # Molecules |
|---|---|---|
| **Cell Death and Survival** | 1.42E-21 - 1.51E-128 | 1649 |
| **Cellular Assembly and Organization** | 1.30E-21 - 6.03E-94 | 1152 |
| **Cellular Function and Maintenance** | 8.71E-22 - 6.03E-94 | 1448 |
| **Protein Synthesis** | 1.77E-22 - 1.34E-89 | 749 |
| **Cell Morphology** | 1.97E-22 - 1.35E-65 | 971 |

#### Physiological System Development and Function

| Name | p-value range | # Molecules |
|---|---|---|
| **Organismal Survival** | 1.53E-99 - 1.53E-99 | 1128 |
| **Tissue Development** | 9.30E-22 - 9.63E-47 | 793 |
| **Cardiovascular System Development and Function** | 8.11E-22 - 8.37E-37 | 690 |
| **Connective Tissue Development and Function** | 3.55E-22 - 1.23E-36 | 491 |
| **Organismal Development** | 2.34E-24 - 4.49E-32 | 1109 |

### Top Tox Functions

#### Assays: Clinical Chemistry and Hematology

| Name | p-value range | # Molecules |
|---|---|---|
| **Increased Levels of Hematocrit** | 4.81E-03 - 4.81E-03 | 25 |
| **Decreased Levels of Albumin** | 1.00E00 - 5.31E-03 | 12 |

(c) 2000-2024 QIAGEN. All rights reserved.

INGENUITY PATHWAY ANALYSIS

| | | |
|---|---|---|
| **Increased Levels of Creatinine** | 1.10E-01 - 2.07E-02 | 18 |
| **Increased Levels of Red Blood Cells** | 3.31E-02 - 3.31E-02 | 25 |
| **Increased Levels of Potassium** | 1.00E00 - 4.78E-02 | 8 |

### Cardiotoxicity

| Name | p-value range | # Molecules |
|---|---|---|
| **Cardiac Enlargement** | 1.00E00 - 6.86E-20 | 245 |
| **Cardiac Necrosis/Cell Death** | 4.68E-01 - 2.01E-12 | 102 |
| **Cardiac Dilation** | 4.68E-01 - 2.01E-11 | 134 |
| **Cardiac Fibrosis** | 6.12E-01 - 7.86E-09 | 91 |
| **Cardiac Arrythmia** | 1.00E00 - 3.53E-07 | 100 |

### Hepatotoxicity

| Name | p-value range | # Molecules |
|---|---|---|
| **Liver Hyperplasia/Hyperproliferation** | 1.00E00 - 1.32E-73 | 1736 |
| **Hepatocellular carcinoma** | 1.00E00 - 2.09E-25 | 529 |
| **Liver Steatosis** | 6.12E-01 - 9.93E-09 | 135 |
| **Liver Necrosis/Cell Death** | 5.46E-01 - 1.23E-08 | 101 |
| **Liver Fibrosis** | 1.00E00 - 6.52E-06 | 125 |

### Nephrotoxicity

(c) 2000-2024 QIAGEN. All rights reserved.

INGENUITY PATHWAY ANALYSIS

| Name | p-value range | # Molecules |
|---|---|---|
| **Renal Necrosis/Cell Death** | 1.00E00 - 2.47E-17 | 211 |
| **Glomerular Injury** | 1.00E00 - 7.27E-09 | 142 |
| **Renal Damage** | 5.06E-01 - 4.94E-08 | 97 |
| **Renal Tubule Injury** | 5.06E-01 - 4.94E-08 | 47 |
| **Renal Proliferation** | 5.46E-01 - 7.11E-07 | 91 |

### Top Regulator Effect Networks

| ID | Regulators | Disease & Functions | Consistency Score |
|---|---|---|---|
| **1** | MLXIPL | Cell death of osteosarcoma cells | 6.0 |
| **2** | RICTOR | Cell death of osteosarcoma cells | 6.0 |
| **3** | LARP1 | Cell death of osteosarcoma cells | 5.833 |
| **4** | MYCN | Cell death of osteosarcoma cells | 5.388 |
| **5** | Lh | Cell death of osteosarcoma cells | 5.196 |

### Top Networks

| ID | Associated Network Functions | Score |
|---|---|---|
| **1** | Cancer, Organismal Injury and Abnormalities, Cardiovascular Disease | 34 |
| **2** | Developmental Disorder, Hereditary Disorder, Metabolic Disease | 34 |

(c) 2000-2024 QIAGEN. All rights reserved.

| | | |
|---|---|---|
| 3 | Hereditary Disorder, Organismal Injury and Abnormalities, Skeletal and Muscular Disorders | 34 |
| 4 | Hereditary Disorder, Neurological Disease, Organismal Injury and Abnormalities | 34 |
| 5 | Cellular Assembly and Organization, Metabolic Disease, Organismal Injury and Abnormalities | 32 |

### Top Tox Lists

| Name | p-value | Overlap |
|---|---|---|
| Renal Necrosis/Cell Death | 1.09E-31 | 32.6 % 211/648 |
| Mitochondrial Dysfunction | 1.43E-27 | 49.7 % 85/171 |
| NRF2-mediated Oxidative Stress Response | 1.42E-16 | 36.0 % 86/239 |
| Cardiac Necrosis/Cell Death | 3.03E-14 | 31.0 % 101/326 |
| Liver Necrosis/Cell Death | 6.81E-13 | 29.6 % 101/341 |

### Top My Lists

### Top My Pathways

(c) 2000-2024 QIAGEN. All rights reserved.

INGENUITY PATHWAY ANALYSIS

### Top ML Disease Pathways

### Top Analysis-Ready Molecules

#### Expr Other

| Molecules | Expr. Value | Chart |
|---|---|---|
| VIM | ↑ 10.722 | |
| ACTG1 | ↑ 10.621 | |
| AHNAK | ↑ 10.423 | |
| MYH9 | ↑ 10.167 | |
| FN1 | ↑ 10.157 | |
| FLNA | ↑ 9.752 | |
| PLEC | ↑ 9.594 | |
| EEF1A1 | ↑ 9.159 | |
| COL1A1 | ↑ 9.100 | |
| TNC | ↑ 9.084 | |

#### Expr Other

| Molecules | Expr. Value | Chart |
|---|---|---|
| ZZEF1 | ↓ -0.585 | |
| ZNF207 | ↓ -0.585 | |
| ZDHHC5 | ↓ -0.585 | |
| YIPF5 | ↓ -0.585 | |

(c) 2000-2024 QIAGEN. All rights reserved.

INGENUITY PATHWAY ANALYSIS

| | | |
|---|---|---|
| **XPO5** | ↓ | -0.585 |
| **XPNPEP3** | ↓ | -0.585 |
| **WNK1** | ↓ | -0.585 |
| **WDR36** | ↓ | -0.585 |
| **VPS26B** | ↓ | -0.585 |
| **VCPIP1** | ↓ | -0.585 |

(c) 2000-2024 QIAGEN. All rights reserved.

INGENUITY PATHWAY ANALYSIS
